# CRISPRi screen for enhancing heterologous *α*-amylase yield in *Bacillus subtilis*

**DOI:** 10.1101/2022.03.30.486407

**Authors:** Adrian Sven Geissler, Annaleigh Ohrt Fehler, Line Dahl Poulsen, Enrique González-Tortuero, Thomas Beuchert Kallehauge, Ferhat Alkan, Christian Anthon, Stefan Ernst Seemann, Michael Dolberg Rasmussen, Anne Breüner, Carsten Hjort, Jeppe Vinther, Jan Gorodkin

**Affiliations:** Center for non-coding RNA in Technology and Health, Department of Veterinary and Animal Sciences, University of Copenhagen, Denmark; Section for Computational and RNA Biology, Department of Biology, University of Copenhagen, Denmark; Novozymes A/S, Denmark

## Abstract

Enhancing yield during bacterial enzyme production could have positive economic and environmental impacts. For cell factories, such improvements in yields could potentially be obtained by fine-tuning the metabolic processes and their regulatory mechanisms for gene candidates. In pursuit of such candidates, we performed RNA-sequencing of two *α*-amylase producing *Bacillus* strains and predict hundreds of putative novel non-coding transcribed regions. Complex operons that are regulated by a wide variety of transcription factors, non-coding and structured RNAs add to the challenge of finding yield-affecting candidates. Surprisingly, we found that non-coding genomic regions are proportionally undergoing the highest changes in expression during fermentation (75% of novel RNA predictions had absolute logFC > 2). Since these classes of RNA are also understudied, we targeted the corresponding genomic regions with CRIPSRi knockdown to test for any potential impact on the yield. From differentially expressed annotations, including both novel candidate and prior annotated ncRNAs, we selected 53 non-coding candidates. The targeting with CRISPRi knockdowns transcription in a genomic region on both the sense and the antisense strand. Thus, the CRISPRi experiment cannot link causes for yield changes to the sense or antisense disruption. Nevertheless, we observed on several instances with strong changes in enzyme yield. The knockdown targeting the genomic region for a putative antisense RNA of the 3’ UTR of the *skfA-skfH* operon led to a 21% increase in yield. In contrast, the knockdown targeting the genomic regions of putative antisense RNAs of the cytochrome *c* oxidase subunit 1 (*ctaD*), the sigma factor *sigH*, and the uncharacterized gene *yhfT* decreased yields by 31 to 43%.

## Introduction

*α*-amylases are essential enzymes in commercial applications, representing about 25-33% of the world enzyme market (Nguyen et al., 2002). They are used in various applications, such as in detergents and in the paper, leather, and pharmaceutical industries (de Souza & Magalhães, 2010). Therefore, improving the efficiency of *α*-amylase production would affect a broad range of industries and have a beneficial economic and environmental impact. Although various *Bacillus* species can industrially produce *α*-amylases (Schallmey et al., 2004), this study focuses on *B. subtilis* due to its high biotechnological versatility (Hohmann et al., 2016; van Dijl & Hecker, 2013). Various genetic modifications can achieve commercial-scale production yields in *Bacillus* organisms. These modifications commonly optimize the protein secretion system (Kontinen & Sarvas, 1993; Quesada-Ganuza et al., 2019; Vitikainen et al., 2001), metabolic pathways (Fischer & Sauer, 2005), or signaling pathways (Davidson et al., 2012; Hohmann et al., 2016; Veening et al., 2008). Notably, the yields increase with an optimization of the *α*-amylase’s gene sequence. A higher *α*-amylase expression and thus higher yield can be achieved by increasing the strength of the promoter (Hohmann et al., 2016), substitution of rare codons (Quax et al., 2015), and destabilization of mRNA secondary structures (Kudla et al., 2009). Overall, these improvement approaches share a focus on protein-coding genes and, to some extent, their RNA structures. However, the broader class of regulatory non-coding genes and RNA structures (ncRNA) are relatively unexplored in the context of enzyme production optimization (Hohmann et al., 2016). Nevertheless, experiments from a *B. licheniformis* production strain producing alkaline serine protease found an abundance of non-coding RNA elements expressed during fermentation (Wiegand, Dietrich, et al., 2013; Wiegand, Voigt, et al., 2013). Thus, finding potential yield improving candidates remains challenging, particularly when also considering the co-transcribed genes of bacterial polycistronic operons.

While finding potential candidates is already challenging, testing and assessing their impact on enzyme yields is an additional challenge: Traditional knockout strains with subsequent yield assessment is resource-intensive and demand preceding ranking of candidates based on computational analysis to improve the hit rate (Hohmann et al., 2016). One approach is to predict changes in yield by simulating the complex expression dynamics of regulatory mechanism and the flow in metabolic pathways (Hohmann et al., 2016). Such predictions depend on detailed models of metabolic pathways and regulatory interactions (Caspi et al., 2014; M. Kanehisa, 2000; King et al., 2016; Szklarczyk et al., 2019). Further, the creation of such models is a non-trivial endeavor (King et al., 2016; Szklarczyk et al., 2019), especially for ncRNAs (Buescher et al., 2012). For ncRNAs without prior knowledge of functions and regulatory interactions, this type of modeling is impossible without additional experimental information. Potential regulatory associations can be identified by analyzing expression patterns for a number of experimental conditions (Huynh-Thu et al., 2010; Leong et al., 2014; Nicolas et al., 2012; Nolte et al., 2019) with subsequent filtering for regulatory interactions using statistical methods (Huynh-Thu et al., 2010; Leong et al., 2014; Nolte et al., 2019).

Here, our goal is to target genomic regions (spanning both coding and non-coding genes) that potentially impact enzyme yield. We are not concerned about mechanisms of specific genes and do therefore employ strand-unspecific search for such genomic regions. For example, differential expression of an antisense RNA (asRNA) can point to its genomic region as a yield-changing candidate, but whether this asRNA or the sense located operon is the cause or effect is here beyond the scope. To obtain the genomic candidate regions, we conducted RNA-seq experiments to identify expressed genes during fed-batch fermentation of *Bacillus subtilis* strains, genetically manipulated to over-express *α*-amylase. These regions then targeted with CRISPRi to test the effect on yield of the α-amylase enzyme.

## Materials and Methods

### Strains and media

*B. subtilis* strain 168 Δ*spoIIAC* Δ*amyE*::*spec-*P_4199_-*prsA* Δ*apr* Δ*nprE* Δ*srfAC* was maintained on LBPG medium at room temperature. Two different strains were generated, one with a codon-optimized α-amylase JE1 sequence (*prsA+je1zyn*) and one with the wild-type sequence (*prsA+je1*) with the genomic annotation Δ*pel::P4199-je1-cat* and Δ*pel::P4199-je1zyn-cat*, respectively. The DNA sequence of the inserted fragments was verified in both strains. A pairwise sequence alignment of the strains’ genomic sequences against the reference assembly (see “Library preparation and sequencing”) confirms the above listed mutagenic deletions and insertions. However, the alignment shows that the ICEBs1 mobile element (Auchtung et al., 2007; Johnson & Grossman, 2015) has been lost in the strains used in this study, since a gap is observed exactly between the two attachment sites *attL* and *attR* (C. A. Lee et al., 2007). The excision event has somehow retained a 240 bp stretch of the gene *cwlT* (Johnson & Grossman, 2015). Aside of deletions of 55 bp in *rpoE, 47 bp* in *rrnH-16S, and104 bp* in *rrnH-23S,* none of which are essential genes, the strains did not contain any genomic structural variations relative to the reference genome. If not indicated otherwise, strains were cultivated at 37 °C in tryptone-yeast (TY) medium supplemented with 1 µg/ml erythromycin when appropriate.

For the bioreactor fermentations, as a first inoculum medium, the strains grew in SSB4 agar, which is a complex medium containing, per liter of deionized water: Soy peptone SE50MK (DMV) 10 g, sucrose 10 g, Na_2_HPO_4_ *×*2H_2_O 5 g, KH_2_PO_4_ 2 g, citric acid 0.2 g, thiamin hydrochloride 11.4 mg, calcium D-pantothenate 9.5 mg, nicotinic amide 7.8 mg, folic acid 2.9 mg, pyridoxal-HCI 1.9 mg, riboflavin 0.95 mg, D-biotin 0.38 mg, FeSO_4_ *×*7H_2_O 39.3 mg, MnSO_4_ *×*H_2_O 9.8 mg, ZnSO_4_ *×*7H_2_O 8.2 mg, CuSO_4_ *×*5H_2_O 3.9 mg, and agar 25 g. pH was adjusted to 7.3-7.4 with NaOH. Afterward, the transfer buffer was M-9 medium, a chemically defined buffer containing, per liter of deionized water: Na_2_HPO_4_ *×*2H_2_O 8.8 g, NaCl 4 g, KH_2_PO_4_ 3 g, and MgSO_4_ *×*H_2_O 0.2 g. Shake flasks contained PRK-50 medium, which is a complex medium consisting of, per liter, soy grits 110 g, and Na_2_HPO_4_ *×*2H_2_O 5 g. pH was adjusted to 8.0 with NaOH (4N, 13,9% w/v) / H3PO4 (16% v/w) before sterilization. The make-up medium was a complex medium composed by, per liter, tryptone (Difco) 30 g, KH_2_PO_4_ 7 g, Na_2_HPO_4_ *×*2H_2_O 7 g, K_2_SO_4_ 5 g, MgSO_4_ *×*7H_2_O 4 g, (NH_4_)_2_SO_4_ 4 g, citric acid 0.78 g, thiamin hydrochloride 34.2 mg, calcium D-pantothenate 28.4 mg, nicotinic amide 23.3 mg, pyridoxal-HCI 5.7 mg, riboflavin 2.8 mg, folic acid 2.5 mg, D-biotin 1.1 mg, FeSO_4_ *×*7H_2_O 157 mg, MnSO_4_ *×*H_2_O 39.2 mg, ZnSO_4_ *×*7H_2_O 32.8 mg, CuSO_4_ *×*5H_2_O 15.6 mg, and antifoam (SB2121) 1.25 ml. pH was adjusted to 6.0 with NaOH (4N, 13,9% w/v) / H3PO4 (16% v/w) before sterilization. Sucrose 2M was employed as a feed medium.

### Construction of *B. subtilis* strains containing heterologous genes

Splicing by Overlapping Extension-PCR (SOE-PCR) method (Horton et al., 1989) was used to generate linear recombinant DNA for transformation. The *α*-amylase JE1 was obtained from *Bacillus halmapalus,* and it was later codon-optimized for *B. subtilis* into je1zyn with a Novozymes proprietary codon optimization model. Synthetic genes encoding *je1, je1zyn,* and dCas9 were ordered from GeneArt. Finally, an additional chaperone PrsA was added to increase the secretion performance of the Sec translocation pathway (Hohmann et al., 2016; van Dijl & Hecker, 2013). Recombinant DNA was directed to a specific locus by the addition of flanking regions containing sequences homologous to that locus. An antibiotic resistance marker gene (either chloramphenicol [*cat*] or spectinomycin [*specR*]) was also included to enable selection for strains with the fragments integrated in the chromosome. Overexpression of sRNAs was driven by the P4199 promoter (Joergensen, 1999). The sRNA genes were amplified from genomic DNA from *B. subtilis* str. 168 and joined to flanking regions enabling integration into the *alr* locus as described above. Clones in which a double cross-over event had occurred were selected for on LB agar plates containing the appropriate antibiotic. The final clones were verified by Sanger sequencing.

### Bioreactor fermentation

The fermentations were conducted in duplicates in custom Novozymes-made 5L tanks. First, both strains (*prsA+je1* and *prsA+je1zyn*) were grown on SSB-4 agar slants 1 day at 37 °C separately. The agar was then washed with M-9 buffer, and the optical density at 650 nm (OD_650_) of the resulting cell suspension was measured to calculate the number of cells. Shake flasks containing PRK-50 medium were incubated at 37 °C at 300 rpm for 20 hours with an inoculum of OD_650_ reaching 0.1 (equivalent to 10^8^ CFU/ml). Fermentation in the tank was started by inoculating it with the growing culture from the shake flask. The inoculated volume made up 11% of the total medium volume (80 ml for 720 ml fermentation medium).

Standard lab fermentations were performed at 38 °C (controlled by a temperature control system), with a pH of 6.8 – 7.2 (regulated with NH_4_OH and H_3_PO_4_, respectively), aeration of 1.5 l/min/kg broth weight, and agitation of 1500 rpm. The feed strategy started with a 0.05 g/min/kg initial broth after inoculation (0 hours) and shifted to 0.156 g/min/kg initial broth after inoculation until the end. The cultivation was run for five days with constant agitation, and the oxygen tension was measured with a dissolved oxygen electrode and followed on-line in this period. The different strains were compared side by side, and OD_650_ measurements were done to calculate the number of bacterial cells in the fermentation. Finally, JE1 amylase activities for in culture supernatants diluted to 1/6000 in Stabilizer buffer were measured with an in-house activity measure in [KNU(N)/g]. KNU stands for Kilo Novo *α*-amylase Units of Natalase (commercial name for JE1). The activity KNU is the amount of enzyme which breaks down 5.26 g starch per hour. The overall activity is normalized by the amount of starch in grams used in the activity assay.

### High-throughput sgRNA cloning

The Pq promoter was used to express the sgRNA. The first expression cassettes (including the Pq promoter, sgRNA target sequence, sgRNA constant domain, and terminator) were ordered from GeneArt as a DNA string with a sgRNA target sequence directed towards GFP (5’-TCTGTTAGTGGAGAGGGTGA-3’, pTK0001) and the signal peptide sequence for *je1* and *je1zyn* (5’-GAATCATGAAACAACAAAAA-3’, the same signal peptide sequence used for both wild type and codon-optimized constructs, pTK0002) (see sequences in Supplementary file 4). The sgRNA expression cassette was cloned into pE194 by POE PCR (You & Percival Zhang, 2014). Transformants were plated on erythromycin (1 µg/mL) LBPG medium and positive clones were identified by colony PCR.

To integrate the sgRNA expression construct, the episomally cloned sgRNA::GFP expression cassette was moved into the *alr* locus of *B. subtilis* strain 168 Δ*spoIIAC* Δ*amyE* Δ*apr* Δ*nprE* Δ*srfAC pel*::P4199-*je1zyn*-*cat*, *amyE*::*spec* P4199’ *dcas9* insert. The resulting strain carried a disruption in the *alr* gene, resulting in a D-alanine auxotroph strain. Flanking sequences, which direct the homologous recombination, were amplified from a wild type strain carrying the functional *alr*. These regions were combined with the sgRNA::GFP expression cassette amplified from pTK0001 by SOE. Upon successful transformation and integration of the final SOE product, the disrupted *alr* gene was repaired. The transformation was performed as previously described in (Yasbin et al., 1975), with the addition of 400 µL 10 mg/mL D-alanine to 10 ml Spitz transformation media (Sadaie & Kada, 1983).

For the HTP cloning of sgRNA, the 20bp sgRNA target sequence in the chromosomally integrated sgRNA::GFP was substituted with new 20 bp target sequences by oligo overlap (complete oligo cloning sequences in table Supplementary file 3). Briefly, the new sgRNA target sequence oligos (up_sgRNA and down_sgRNA) were ordered in plates from Eurofins Genomics. The up_sgRNA oligo was combined with pep0945 to create the up_sgRNA_fragment (Ap), and the down_sgRNA oligo was combined with oTK0274 to generate the down_sgRNA_fragment (Bp). Template Ap and Bp were then combined, and together with pep0946 and pep0948, the new SOE fragment carrying the new sgRNA sequence was created. The SOE was transformed in *B. subtilis* strain 168 Δ*spoIIAC* Δ*amyE* Δ*apr* Δ*nprE* Δ*srfAC pel*::P4199-*je1zyn*-*cat*, *amyE*::*spec* P4199’ *dcas9* insert as described above. The final clone was verified by Sanger sequencing.

### Deep-well plate fermentation

The System Duetz system (Enzyscreen), consisting of sandwich covers (Enzyscreen #CR1996) and 96-well deep-well plates (Enzyscreen #CR1496b) combined with the corresponding clamp system (Enzyscreen #CR1700), was used for small scale cultivations. Strains were grown in 500 µl TY medium at 37 °C, 300 rpm with a 2-inch throw radius. Plates were inoculated from cryo-stocks using the Cryo-replicator press (Enzyscreen #CR1100) and grown overnight. A fresh plate was inoculated with 10 µl culture and grown at 37 °C, 300 rpm for 24 h. The supernatant for amylase activity measurements was harvested by centrifugation at 4000 rpm for 10 min. Optical density at 450 nm (OD_450_) was measured to estimate the bacterial growth curve using (VWR V-3000PC spectrophotometer). The final OD_450_ values were determined by subtracting the 2.51-multiplied sample well value with the blank well value.

Additionally, JE1 amylase activities were measured in culture supernatants using the AMYL kit (Roche/Hitachi #11876473 001, note: this assay kit has recently been discontinued). In this case, the culture supernatants were diluted to 1/50 in Stabilizer buffer (CaCl_2_ 0.03M and Brij 35 0.0083% (w/v)). Reagent 1 and reagent 2 of the AMYL kit were mixed 10:1 to generate the assay substrate. 20 µl diluted sample was mixed with 180 µl assay substrate. The assay was incubated at 37 °C with shaking for 30 min. Absorbance, defined as the optical density at 405 nm (OD_450_), was measured in a plate reader (Molecular devices Spectramax 340PC). The JE1 enzyme activities were measured in KNU per gram as described for the bioreactor fermentation, although no additional dilution was needed. Due to the number of candidates (see “Retrieval of the candidate list for yield change assessment”), the deep-well plate experiment was run in two batches. The sgRNAs against the “novel asRNA for ybgB” were added in both batches showing similar mean normalized yield impact (paired two-sided t-test p≥0.18).

### RNA purification

Samples from fermentations were harvested by mixing 5 ml cell culture with 5 ml 100% ethanol, immediately storing on dry ice before transferring to −80 °C. Cells were pelleted by centrifugation for 5 min at 1780 g at −9°C. Samples for qRT-PCR was obtained from triplicate overnight cultures that were diluted to OD_450_ 0.05 before harvesting 10 ml culture at OD_450_ ∼ 0.8 on ice and cells were immediately collected at 3,220 g for 4 min at 4 °C. All pellets were vortexed for 4 min in 1 ml RNA extraction buffer (80 mM LiCl, 8 mM EDTA, 8 mM Tris-HCl (pH 7.4), 0.2% SDS, 250 mM NaOAc (pH 4), 8 mM MgCl2 for fermentation samples and 10 mM NaOAc, 150 mM sucrose, 1% SDS for qRT-PCR samples), 1 ml phenol:chloroform 5:1 pH 4.5 (Thermofisher #AM9720) and 0.5 ml glass beads (Sigma #G8772). Samples for qRT-PCR were incubated for 5 min at 65 °C before freezing in liquid nitrogen. All samples were then centrifuged for 20 min at 17,000 g at 4 °C before transferring the aqueous phase to repeat the phenol extraction. The aqueous phase was then mixed with 1 volume of chloroform before centrifugation at 13,000 g for 10 min at 4 °C for phase separation. RNA was finally precipitated in 1 volume isopropanol at room temperature for 10 min before centrifugation at 15,000 g for 45 min at 4 °C. RNA pellets were washed with 70% ethanol and dissolved in water.

DNase digestion of RNA samples for qRT-PCR was performed using TURBO DNase (Invitrogen #AM2238) and purified using RNA Clean & Concentrator (Zymo research #R1016) before assessing RNA integrity using gel electrophoresis. For fermentation RNA samples, DNase treatment was carried out in solution using the RNase-free DNase Set (Qiagen #79254) and purified with the RNeasy MinElute spin columns (Qiagen #74204) before assessing RNA integrity on bioanalyzer.

### Quantitative RT-PCR

RNA was obtained from exponentially growing cultures in TY media (see “Strains and media” and “RNA purification” above). Quantitative RT-PCR was performed using Brilliant III Ultra-Fast SYBR Green qRT-PCR Master Mix (Agilent Technologies #600886) according to manufacturer’s protocol with 5 ng RNA in 10 µl reactions using 0.5 µM of each primer (Supplementary file 4). Each of the three biological replicates were quantified in technical duplicates using Quantstudio 6 Flex (Applied Biosystems #4485694) incubating at 50°C for 10 min, 95 °C for 3 min and 40 cycles of 95 °C for 5 seconds and 60 °C for 15 seconds. Fold changes were calculated using the 2^-ΔΔCt^ method and citA was used as reference gene.

### Library preparation and sequencing

Bacterial RNA library preparation was prepared from rRNA-depleted total RNA as previously published (Poulsen & Vinther, 2018). The library was reverse transcribed with a random priming position. The libraries were single-end sequenced by the MOMA NGS Core Center at the Aarhus University Hospital, Denmark, on an Illumina NextSeq 500. The read lengths were 76 bp; for details on primer, multiplexing indices, and adapter sequences, see section S1 in Supplementary file 1. The quality of the sequenced reads was assessed with FastQC v. 0.11.8 (S. Andrews, 2018). Then, to remove adapter contamination from the reads, trimmomatic tool v. 0.39 (Bolger et al., 2014) was run, allowing two seed mismatches, clip sequences of at least 10 bp overlaps, and, at least, 30 bp in case of palindromic overlaps. A 4-bp sliding window size was set to ensure an average quality above a PHRED score of 20. Trialing bases with a quality score below 3 were removed, and only fragments of at least 40 bp length were kept. Trimmomatic was also used to remove the PCR random index sequence after reducing PCR bias with the clumpify/de-duplication feature of BBMap v. 38.69 (Bushnell et al., 2017). These cleaned reads were mapped against the genome sequences of the respective strains with segemehl v. 0.3.4 (Pedregosa et al., 2011) with the E-value cut-off for seeds set to 5 and overall minimal accuracy of the semi-global alignment of 95%. The resulting statistics of the read filtering and mapping are in Table S1.2.

For optimal reproducibility, all described computational analyses (including the subsequent ncRNA prediction and expression analysis) were compiled in a snakemake v. 5.7.1 workflow (Köster & Rahmann, 2012) and nested in a conda environment v. 4.7.12 (Fig. S1.3). If not otherwise stated, genome reference annotations are according to the most recent comprehensive transcript and non-coding gene annotation of the *Bacillus subtilis* genome (BSGatlas v.1.1, http://rth.dk/resources/bsgatlas (Geissler et al., 2021)).

### Prediction of novel ncRNAs

In order to better cover the complexity of transcriptional regulation and bacterial operons (see Introduction), we complement the existing gene annotation with predicted ncRNAs from the RNA-seq data. Although the genomic sequences of the two substrains and the reference genome used in the BSGatlas (Geissler et al., 2021) differ slightly, the coordinates were matched with *liftOver* (Haeussler et al., 2019) from pairwise alignments with LASTZ (Harris, 2007). Based on the transferred transcription coverages, both for multi-mapping and uniquely mapping reads separately; transcribed regions were computed with the transcript prediction feature of ANNOgesic v. 1.0.8 (Yu et al., 2018). The tool was benchmarked for all parameter combinations consisting of (i) the coverage height-cutoff 1 through 20 and (ii) the minimal number of required replicates ranging 1 to 4 (for details see S2). The quality of the predicted transcripts was assessed by comparing these to the set of all known operons, including isoform transcripts and genes without known transcript according to the BSGatlas.

After comparing all overlapping pairs, only those transcripts with a height cut-off of 5 observed in at least two replicates were considered as predicted transcripts. Under usage of the genome annotation utility *plyranges* v. 1.2.0 (S. Lee et al., 2019), overlapping predicted transcribed regions from both the multi-mapping and uniquely mapping based predictions were combined. From this the resulting novel transcribed regions (NTR) were extracted via intersection with the annotation gaps in the comparison reference set. NTRs of a minimal length of 50 bp (see length distribution in Fig S2.3B) were selected and *in silico* classified as (i) asRNA if at least 90% of the NTR was overlapped by reference annotation on the opposite strand, (ii) UTR if the nearest reference annotation on the same strand was closer than 100 bp, (iii) novel ncRNA transcribed if the distance was above 1,000 bp, (iv) and as an unclear case if the distance was in-between these two values. (v) The special case of NTR having any antisense overlap with an rRNA gene (without cut-off) was accordingly noted. A full table of the predicted ncRNAs is in Supplementary file 2; the BED format of the predictions is included in Supplementary file 5. The putative classification types are indicated in the BED files by coloration (Fig. S2.5).

### Differential gene expression

The gene expressions of the putative novel ncRNAs found in the preceding step, the coding- and non-coding gene annotations of the BSGatlas, and the strain-specific, synthetic genes for both uniquely mapping and multi-mapping reads were quantified with *featureCounts* v. 1.6.4 (Liao et al., 2014). Read mappings overlapping at least 50% of a gene annotation were considered for quantification, and multi-mapping reads were not weighted. According to the NCBI reference genome sequence, the annotation coordinates were first lifted over to the individual strain sequences before quantification (Fig. S1.3).

Ribosomal RNAs (rRNAs), transfer-messenger RNAs (tmRNAs), and the signal recognition particle (SRP) genes were excluded from the subsequent analysis steps, such that the gene content and raw-expressions between the libraries became comparable (Figs. S3.6 and S3.7 Supplementary file 1). Changes in expressions and the impact on the library size-normalization of *DESeq2* v. 1.22.1 (Love et al., 2014) when including multi-mapping reads were investigated. The size-factor normalization factors did only numerically neglectable change and were overall perfectly correlated (*P* < 2.2 x 10^-16^; Pearson’s product-moment correlation test), and the overall expression values after normalization only increased when considering multi-mapping reads. In contrast to the ncRNA prediction step, the expression analysis was conducted under consideration of multi-mapping reads without an explicit uniquely-mapping only scenario. Gene selection and normalization method was validated with a principal component analysis (PCA) plot on the blind *r*-log-transformed 500 most variable expressed genes, which showed the biological replicates in proximity as anticipated (Fig. 1C).

**Figure 1.**
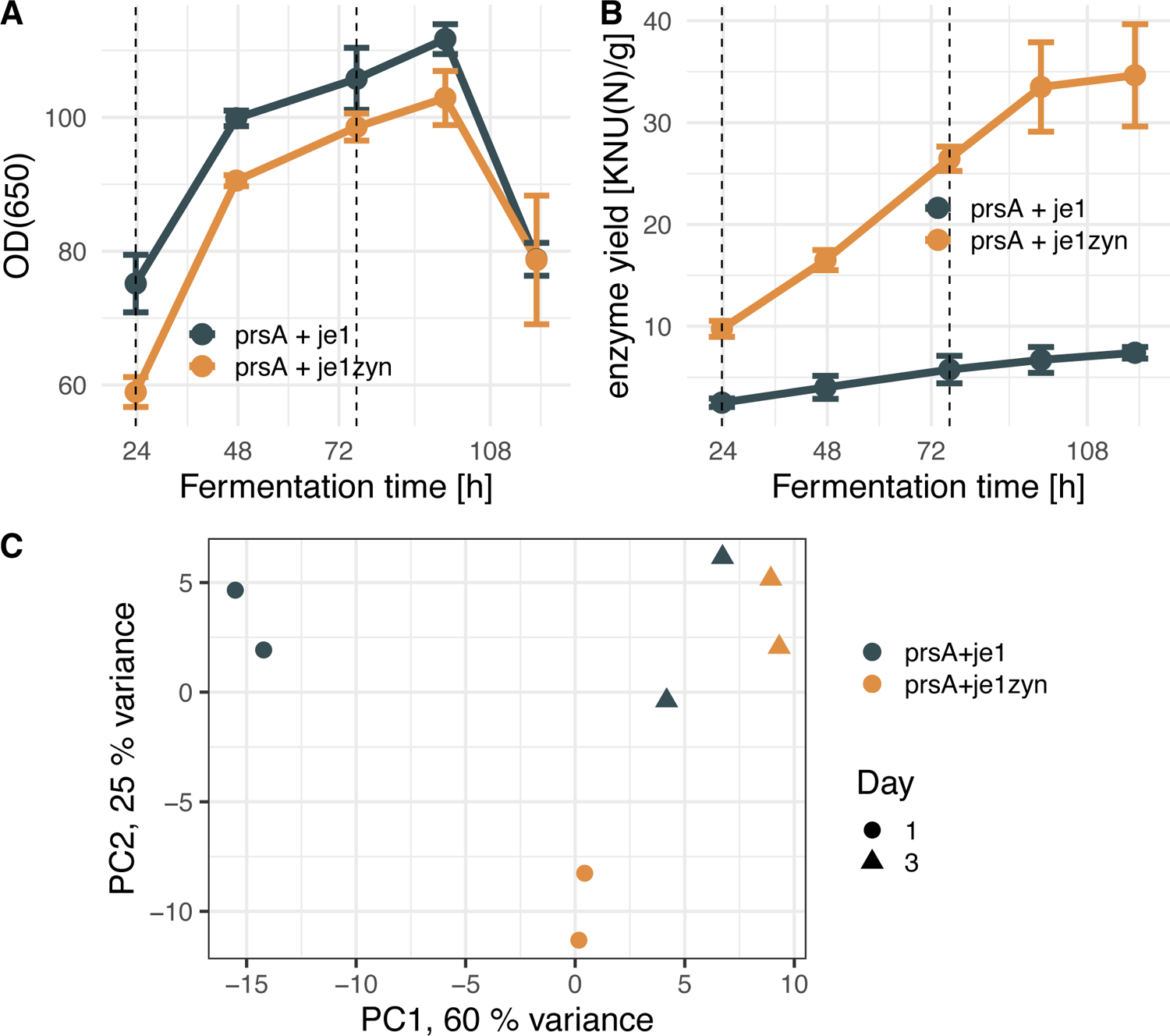
RNA-seq of fermentation samples of *B. subtilis* strains. (A) Fermentation growth curve of a codon optimized strain *prsA+je1zyn* (orange) and control strain *prsA+je1* (grey) (n = 2, error bars depict standard deviation). Time-points for sampling for RNA-seq are indicated with vertical dashed lines. (B) Yield of the *α*-amylase enzyme as in (A). (C) Principal component analysis of RNA-seq samples based on DESeq2 r-log normalized expression of the 500 most variable expressed genes, excluding rRNA, tmRNA, and SRP.

To detect differential expression, *DESeq2* creates for each gene a regression model of the expression in each strain over time. We used different combinations of contrasts first to identify non-/coding genes that had overall non-constant expression (Wald test on the intercept), and afterward, test which pairwise comparison had a differential expression. There were two pairs for comparison over the time-course of fermentation and two pairs between the strains. An additional test investigated the changes between the two measurement time points (time-strain interaction of the regression model). *P-*values computed for the differential expression between both strains on the last day were further corrected with the *fdrtool* v. 1.2.15 (Strimmer, 2008) (Fig. S3.8), and DESeq2’s independent filtering was manually repeated to remove lowly expressed genes from the analysis. The stage-wise multiple-hypotheses-correction procedure, as implemented by *stageR* v. 1.4.0 (Van den Berge et al., 2017), combined the screening step for non-constantly expressed genes with subsequent confirmation of the five pairwise tests. *P-value*s of the screening step were FDR-corrected. Once a non-constant expression was confirmed, at most, two Null Hypotheses of the pairwise test could be true, which allowed for a modified Holm-Procedure correction of the confirmation step (Shaffer, 1986). An overall FDR adjusted P-value ≤ 0.05 was considered. *K*-means clustering of the mean per condition (time+strain) variance regularized expression (DESeq2’s *r-*log transformation) was used to detect significant features. The number of *K*-means clusters was determined as k=12 via the elbow method (Thorndike, 1953).

### Gene-set enrichment

For each cluster, enrichment tests against the GO terms annotated in the BSGatlas were performed with the Elim algorithm implemented in *topGO* v. 2.34.0 (Alexa & Rahnenfuhrer, 2018) with an alpha of 0.01 and a minimal set size of 10. Similarly, enrichment tests against the KEGG pathways were performed using a hypergeometric test implemented in the Fisher’s exact test (R’s fisher.test) function of R. 3.6.0 (R Core Team, 2019) with an alpha of 0.01 and a minimal set size of 10. Pathway annotations for *B. subtilis* were retrieved via the KEGG REST API (https://www.kegg.jp/kegg/rest/keggapi.html) (M. Kanehisa, 2000; Minoru Kanehisa, 2019).

### Retrieval of the candidate list for yield change assessment

Genomic candidate regions for CRISPRi testing were selected as follows. First, candidates were extracted from the differential expression analysis for known and predicted ncRNAs ranked by the highest observed absolute logFC (adjusted P-value ≤ 0.05). This absolute logFC ranked list was further manually filtered by discarding candidate regions that overlap genes or have genes in its neighborhood (nearest operon) that are metabolic (based on KEGG pathways and GO annotations). Then, the ncRNA annotations were inspected to be antisense to coding genes involved in the enriched KEGG pathways. With outset in the highest absolute logFC and impacted pathways in total 50 candidate regions were selected after expert curation. Second, we selected further candidate regions based on the Rfam (Kalvari et al., 2018) screen and other known ncRNA structures from the BSGatlas. This list contained 7 structures with a significant differential expression and an absolute logFC above 1 in at least one time-point. We initially selected candidates relative to BSGatlas v1.0. The v1.1 update indicated 3 initial novel predicted UTR as known UTR. We still include these 3 candidates in Fig. 6A as known UTR, without the “novel” indication. In total we obtain 53 candidate regions we examine by CRISPRi screens.

### Data availability

The data sets generated and analyzed during the current study, including the RNA-seq data and genome sequences, are available in the GEO (accession number GSE179570). Additionally, the computational pipeline is available at doi 10.5281/zenodo.4534403.

## Results

### RNA sequencing during bacterial enzyme production

To search for yield-associated candidates, we performed fed-batch fermentation of two commercially relevant, non-sporulating *Bacillus subtilis* strains. The commercial strains express *α*-amylase (protein name JE1) of *Bacillus halmapalus*. One of the strains has the original gene nucleotide sequence (*je1*) and the other strain has a codon optimized version of the gene (*je1zyn*); both strains express amylase from the strong P4199 promoter. Both strains are co-expressing the PrsA chaperone from an extra copy of the *prsA* gene, also under control of the P4199 promoter to increase amylase production as is common in industrial applications (Jacobs et al., 1993). We refer to the two strains as *prsA+je1zyn* and *prsA+je1*, respectively. We found that culture densities during fermentation were similar for both strains (Fig. 1A), but the *α*-amylase yield in the *prsA+je1zyn* strain was substantially higher than in *prsA+je1* (Fig. 1B) due to the codon optimization of the je1zyn-gene (*je1* and *je1zyn* encode the same protein JE1). To characterize the transcriptome of the strains, duplicate fermentations were run for each strain and samples were taken out for RNA-preparation after 23.7 h (day 1) and 76.2 h (day 3) because yield and OD increased substantially after 24 h at both time-points. The samples were rRNA depleted and sequenced on an Illumina NextSeq platform, which provided up to 29 Mio. raw reads per library (Table S1.2). All reads were subsequently filtered, processed, and mapped against the strains’ respective genome sequences using an in-house pipeline (see methods). A PCA revealed that the samples group by the experimental parameters strain and day as expected, though the two strains diverge slightly on day 1 (Fig. 1C).

### Novel transcribed regions during fed-batch fermentation

To increase the pool of candidate regions that might impact the yield, we extracted potential novel transcribed regions (NTRs) from the mapped RNA-seq data (see Methods section). We predicted transcribed regions based on sequencing coverage (Yu et al., 2018). By subtracting from transcribed regions those regions with known annotations, we inferred the NTRs (Fig2). Using known gene and transcript annotations as a reference, we identified an optimal set of parameters to determine transcribed regions (Fig. 2B and Supplementary file 1 section S2). We selected a set of balanced parameters with respect to the sensitivity in the number of novel predictions and the recall of existing transcripts versus strength in expression evidence. For the determined parameters, 63% of the known transcript were recalled and the comparison against known transcribed regions indicates that 40% of the predictions were potentially novel (Figure S2.1).

**Figure 2.**
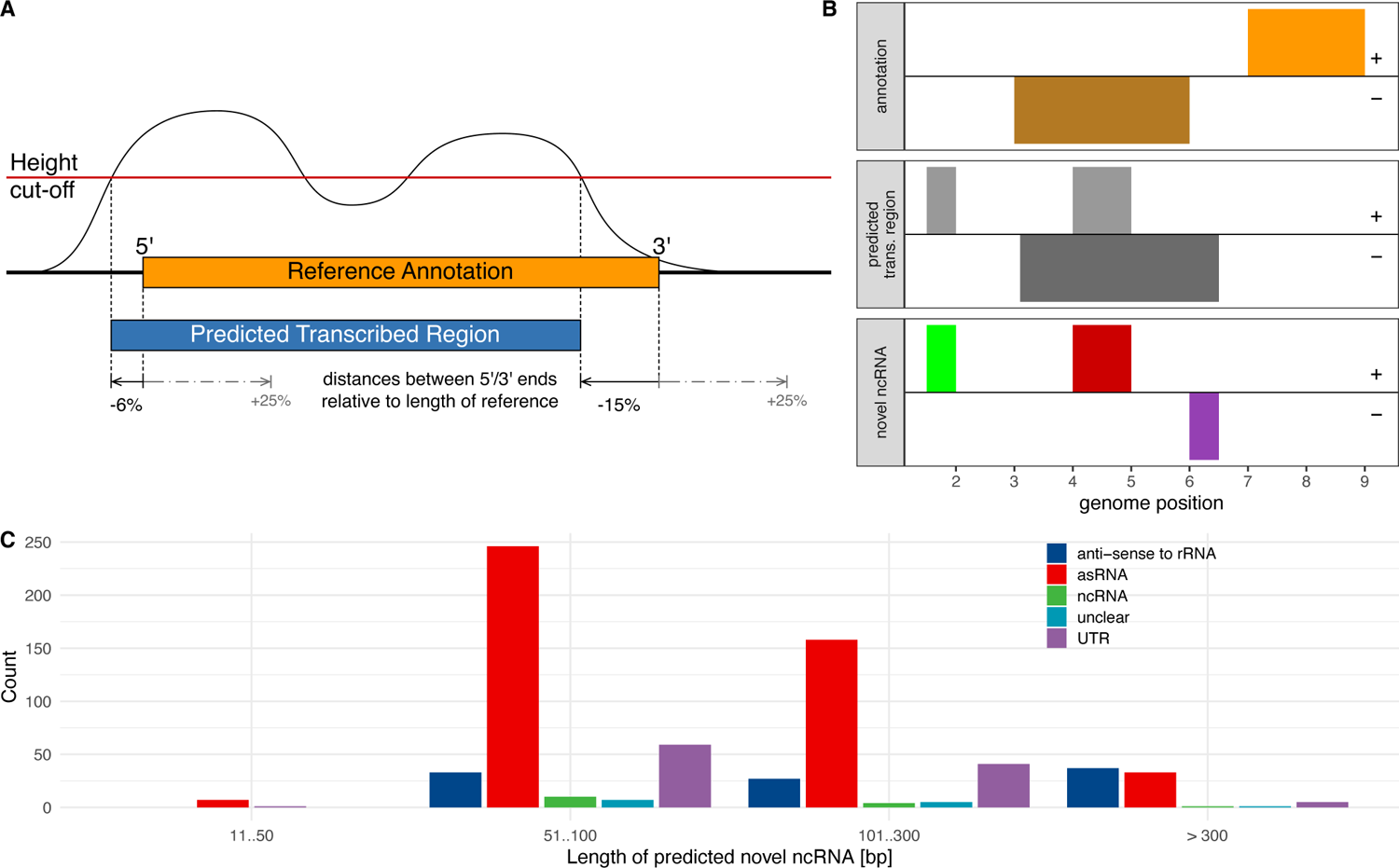
Novel ncRNA prediction. (A) Predicted transcribed regions are regions for which the RNA-seq coverage (black curve) is above a height cut-off (red line). The prediction method tolerated if the coverage for a few base pairs is below the cut-off; this tolerance was larger if these bases were within a reference annotation (see S2 for details). We determined these parameters by benchmarking the transcribed region predictions compared to known reference annotations, particularly their differences in the 5’ and 3’ positions. We measured the predicted transcribed region ends assigned upstream (black arrows) or downstream (grey dashed arrow) of the reference annotation. These values were used in the benchmark (section S2 Supplementary file 1). (B) Novel ncRNAs were identified from parts of transcribed regions (gray) that intersect with strand-specific gaps in the reference annotations (orange). The novel ncRNA predictions are presumably non-coding and depending on the position of these fragments relative to the annotation, they could be classified as novel ncRNAs (green), potentially with antisense overlap (red), or as a UTR (purple). + and – denote strand identity. (C) Number of predicted ncRNAs classification according to length.

After subtracting known annotations, a total of 675 putative novel ncRNA candidates were obtained, which based on their distances to or overlap with known transcript annotations were classified as 106 potential UTRs, 444 asRNA, and 15 potentially independent non-coding transcripts (in the following referred to as potential sRNA) (Fig. S2.3). Many of the novel ncRNA candidates were relatively short (Fig. 2C). In 13 cases, the ncRNAs were classified as both UTR and independent transcripts. Interestingly, there are also 97 ncRNA predictions antisense to rRNA. We did not further specify the potential UTRs as 5’, 3’, or internal UTRs because 56 of the 106 predicted potential UTRs (52.83%) had known annotation close-by (100 nt) both up and down-stream.

### Differential gene expression and pathway analysis

We identified amylase yield changing genomic regions that were differentially expressed during fed batch fermentation by inspecting the expression profiles of the predicted ncRNAs and other known annotations, such as coding genes, non-coding genes, UTRs including cis-regulatory RNA structures (*e.g.,* riboswitches) as annotated in BSGatlas. Using the two time-points and two substrains of the RNA-seq data set, the following five pairwise comparisons were investigated: (i) Differential expression on day 1 and (ii) on day 3 between both strains, (iii + iv) differential expression from day 1 to 3 in either strain, and (v) assessment of the difference between the two fold-changes obtained in i and ii on both days (Fig. S3.4). The naïve approach of testing for differential expression in each case separately and subsequently combining the differentially expressed annotations would cause a substantial loss of statistical power (Van den Berge et al., 2017). Therefore, we conducted a stage-wise hypothesis testing that first screens for dynamic expression before confirming which of the pairwise test applies (section S3 in Supplementary file 1). The procedure guarantees a single overall false discovery rate (FDR) despite that multiple comparisons are investigated for each gene simultaneously (Van den Berge et al., 2017).

At an overall FDR of 5%, the stage-wise testing substantially increases the sensitivity in detecting differentially expressed ncRNA over the naïve approach (Table S3.1). Only five genes with low expression (< 20 per Mio. reads in DESeq2’s base expression value) are detected solely by naïve approach. Stage-wise, on the other hand, detects 966 additional coding (578) and non-coding genes (46) and UTRs (342) to be differentially expressed (Table S3.1). These additional differentially expressed features were highly expressed (more than 100 up to 100,000 per Mio. reads, Fig. S3.2). Of the additionally detected genes, 27 genes were very highly expressed (> 10,000); the five highest expressed genes were the *rny* endoribonuclease (> 100,000), the GTP-binding protein era (> 36,000), the aconitate hydratase *citB* (> 34,000), the sRNA *bsrI* (> 31,000), and the elongation factor *fusA* (> 30,000).

Differentially expressed regions had a wide variety of annotations (Table 1). We observed a differential expression in a total of 1,780 (41.4%) coding genes. We also observed significantly higher transcription of α-amylase on day 1 (logFC > 1.4, p-adj < 1.1e^-6^) and day 3 (logFC > 2.4, p-adj < 5.4e^-16^) for the codon-optimized strain (*prsA+je1zyn*) compared to the non-optimized strain (*prsA+je1*). Differential expression of known non-coding annotations were observed for the RNaseP RNA,11 sRNA (32.4%), 2 asRNA (28.6%), 66 tRNA (76.7%), 50 RNA structures (47.2%, includes riboswitches, cis-regulatory and self-splicing intron structures), and 1,087 of UTRs (19.5%). Of the predicted novel ncRNAs, 372 potential asRNA (65.8%), 13 potential sRNA (72.2%), and 49 potential UTRs (27.2%) were detected as differentially expressed in at least one comparison. The 5 differentially expressed genes having the highest expression (DESeq2’s base mean) were *prsA* encoding the chaperone, the ncRNA srlX with an unknown function, *brsA* encoding 6S sRNA, the JE1 enzyme, and the ribonuclease *rny* (Supplementary file 2). Notably, both *srlX* and *bsrA* were expressed at a higher level than the JE1.

**Table 1.**
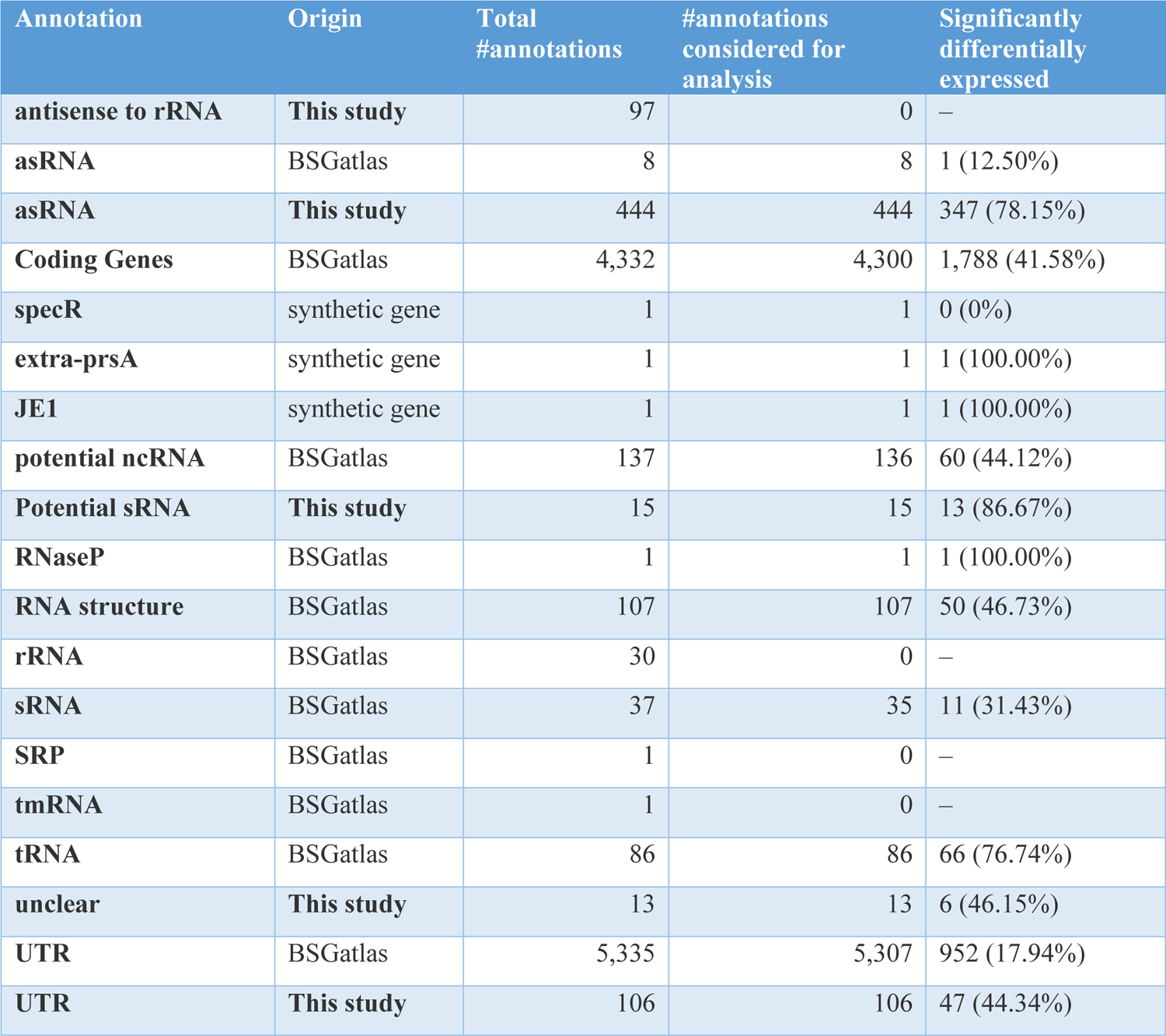
Statistics of differentially expressed genes and other annotations separated by their origin. The possible origins are the predictions from this study, the synthetic genes (see methods), or the BSGatlas. Shown are the total number of annotations and how many of these were considered for differential expression analysis. Not all annotations were considered because of either low overall expression or expression normalization concerns due to deletions relative to the reference genome. Ribosomal RNA, SRP RNA and tmRNA were also excluded from the analysis. The last column indicates how many annotations were significantly differentially expressed in at least one of the tested hypotheses (Fig. S3.4).

Building on the established differential expression analysis procedures to cluster genes with similar expression profiles (Langfelder & Horvath, 2008; Leong et al., 2014; Nicolas et al., 2012; Wiegand, Dietrich, et al., 2013), we associated biological processes through gene-set enrichment tests (Langfelder & Horvath, 2008). Thus, we clustered the genes with similar expression profiles together with k = 12 *k*-means (see methods, Fig. 3 A). Afterward, we conducted gene-set enrichment tests of Gene Ontology (GO) terms (Carbon et al., 2019) and KEGG pathways (Minoru Kanehisa, 2019). 78.3% of all coding genes had GO annotations readily available from the BSGatlas, and pathway annotation could be retrieved for 12.7% of the coding sequences from the KEGG database. Of the 600 available biological processes GO terms, 44 were enriched in at least one *k*-means cluster (topGO with elim term de-correlation, α = 0.01, minimal GO size 10). 10 of 114 KEGG pathways were enriched in an over-representation hypergeometric test (*P* < 0.01, minimal pathway size 10).

**Figure 3.**
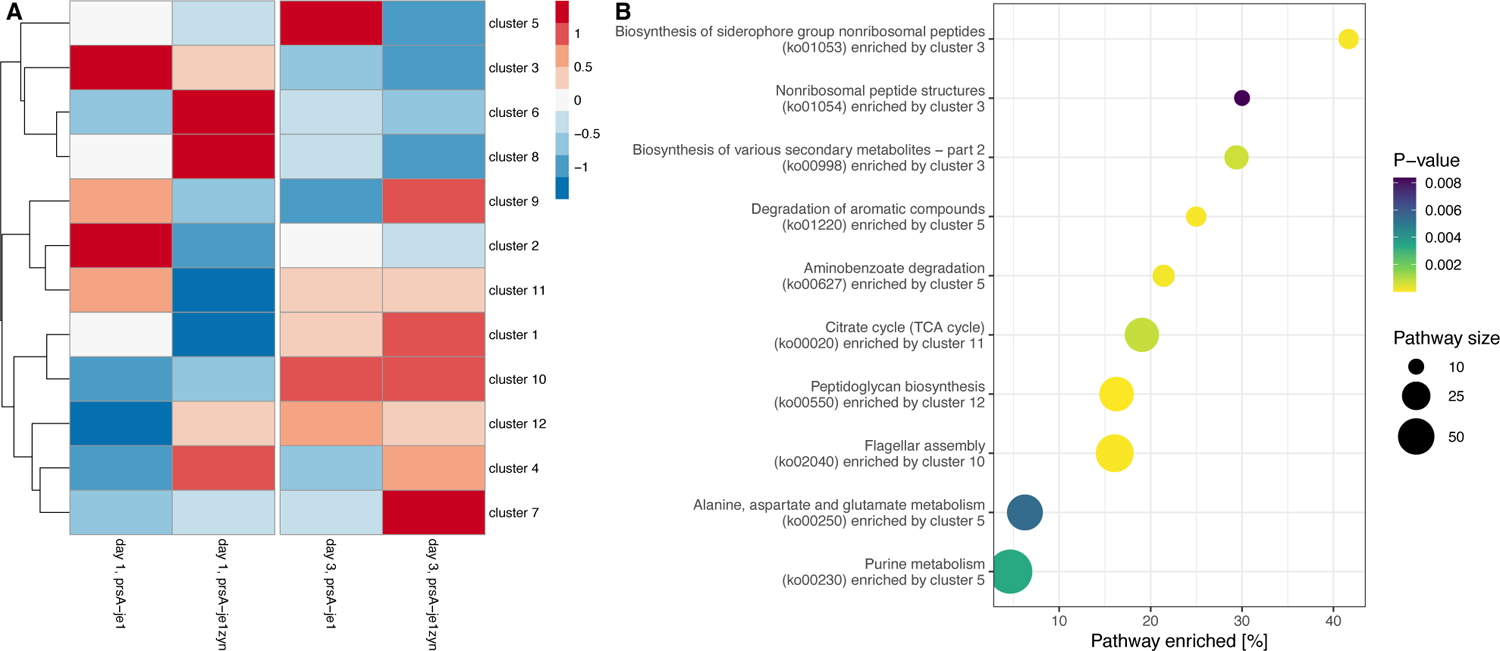
Heatmap and pathway enrichment. (A) Heatmap of the log2 expression profile as implied by the kmeans cluster centroids sorted by a hierarchical complete linkage clustering. The columns contain the expression values at each of the four conditions (days and strain). (B) Enrichment of KEGG pathway by kmeans clusters. The plot shows a pathway and the enriching cluster (y-axis) over the ratio of enrichment (x-axis) with point sizes indicating the number of genes annotated in a pathway and color the P-value of enrichment.

Interestingly, several enriched pathways (Fig. 3 B) are either known targets for optimizing production yields of various metabolites and enzymes or have other molecular relevance. For instance, the central energy citrate cycle (ko00020, cluster 11) and the purine metabolism (ko00230, cluster 5) were enriched. The purine metabolism is a highly regulated pathway and is a known target for biotechnological optimization, having the potential to more than double production yields (Fischer & Sauer, 2005; Hohmann et al., 2016). Some of the enriched pathways are generally relevant for cell growth (biosynthesis of siderophore (ko01053, cluster 3) and peptidoglycan biosynthesis (ko00550, cluster 12)) (S. C. Andrews et al., 2003; Chandrangsu et al., 2017; Vollmer, 2012). The enriched pathways for biosynthesis of various secondary metabolites (ko00998, cluster 3) and non-ribosomal peptide structures (ko01054, cluster 3) are required for the synthesis of antibiotics and sporulation of *B. subtilis* (Demain & Fang, 2000). The enrichment of these two pathways might be a manifestation of the fermentation production stress. The antibiotics survival strategy of *B. subtilis* increases cell motility (Liu et al., 2018), which might explain the enrichment of the flagellar assembly (ko02040, cluster 10). *B. subtilis* fermentation produces side-products such as benzoates and aromatic compounds that cause bacterial stress (Kitko et al., 2009; Singleton et al., 2005); thus, it is not surprising that the corresponding degradation pathways (ko00627, ko01220) were enriched by the same cluster 5. Cluster 5 also enriched the Alanine, aspartate, and glutamate metabolism (ko00250), which further underlines the association with *B. subtilis’* stress response (Feehily & Karatzas, 2013). The GO enrichment indicates the same biological processes that are significantly enriched, although the GO terms provide more detailed information of the involved metabolic processes, stress response, cell motility, and sporulation tendency. The complete list of the enrichment analysis is in Supplementary file 2. Overall, this analysis indicates that clusters of similarly expressed genes enrich pathways that are relevant for enzyme production.

### Yield and differential expression

Given the relevance of our analysis of clusters enriching pathways (last section), the coding and non-coding genes within these clusters are potentially involved in the enriched processes via guilt-by-association (Langfelder & Horvath, 2008). Further, 75% of the predicted asRNA and ncRNA have |logFC| > 2 compared to less than 25% of coding genes (Fig. 5). The expression fold changes (logFC) are significantly greater for novel ncRNA (one-sided Kolmogorov-Smirnov Test p <7.99e-46) and UTRs (p<1.1e-4) compared to coding genes. Additionally, the logFCs observed in coding genes inside operons with antisense located novel asRNA have the tendency of being larger than in all other coding genes (p<1.3e-2). Therefore, we put particular focus on screening for potential yield associations of these novel RNA predictions, in spite that targeting the asRNAs with CRISPRi also disrupts the sense RNA. Based on attributed function of the clusters, their expression, and in some cases known functions from the literature, we selected 53 candidates (32 asRNA, 15 putative ncRNA and 6 known ncRNA, see table in Supplementary file 2). We tested the genomic regions of these candidates for potential effect on production yield. We implemented a CRISPRi framework for knockdown of based on a screening strain stably expressing a catalytically dead Cas9 protein (dCas9). Co-expression of a single guide RNA (sgRNA) allows convenient knockdown of gene expression by recruiting dCas9 to a specific genomic location (Peters et al., 2016). CRISPR-dCas9 targeted strains were grown in the 96 well Duetz system, due to its high reproducibility and throughput (see methods “Deep-well plate fermentation”) (Duetz et al., 2000).

To validate the knockdown of expression levels by dCas9 transcriptional interference, we prepared two strains derived from *prsA+je1zyn*, each expressing a different control sgRNA directed towards *ssrA,* which encodes the transfer-messenger RNA (tmRNA). For the enzyme activity reference point, we normalized activities relative to the effect to sgRNA targeting a plasmid placed *gfp.* Expression of the sgRNAs resulted in a 74 and 98% reduction in expression level of tmRNA, demonstrating efficient CRISPRi knockdown of a ncRNA in our setup (Fig. 4A). The knockdown of the *ssrA* led to a an up to 36% reduced JE1 enzyme activity (Fig. 4B). The tmRNA rescues stalled ribosomes by acting as both a tRNA and mRNA through its short open reading frame which encodes a proteins degradation marker (Karzai et al., 2000). Therefore, the reduction in yield upon tmRNA knockdown implies a potential involvement of the ribosomal rescue mechanism in α-amylase production. Interestingly, the *ssrA* knockdown strain with a more pronounce tmRNA level reduction (Fig. 4A) had a slightly smaller reduction in JE1 yield (< 25%, Fig. 4B). Additionally, we validated the CRISPRi functionality by measuring the GFP fluorescence in the enzyme activity reference strain upon introduction of sgRNA::*gfp* (Fig. 4C). Together, these results show that our CRISPRi system is functional in knockdown both RNA transcript levels and protein abundances. Therefore, the systems allow for an assessment in impact on enzyme yields after knocking down genes in *Bacillus subtilis*.

**Figure 4.**
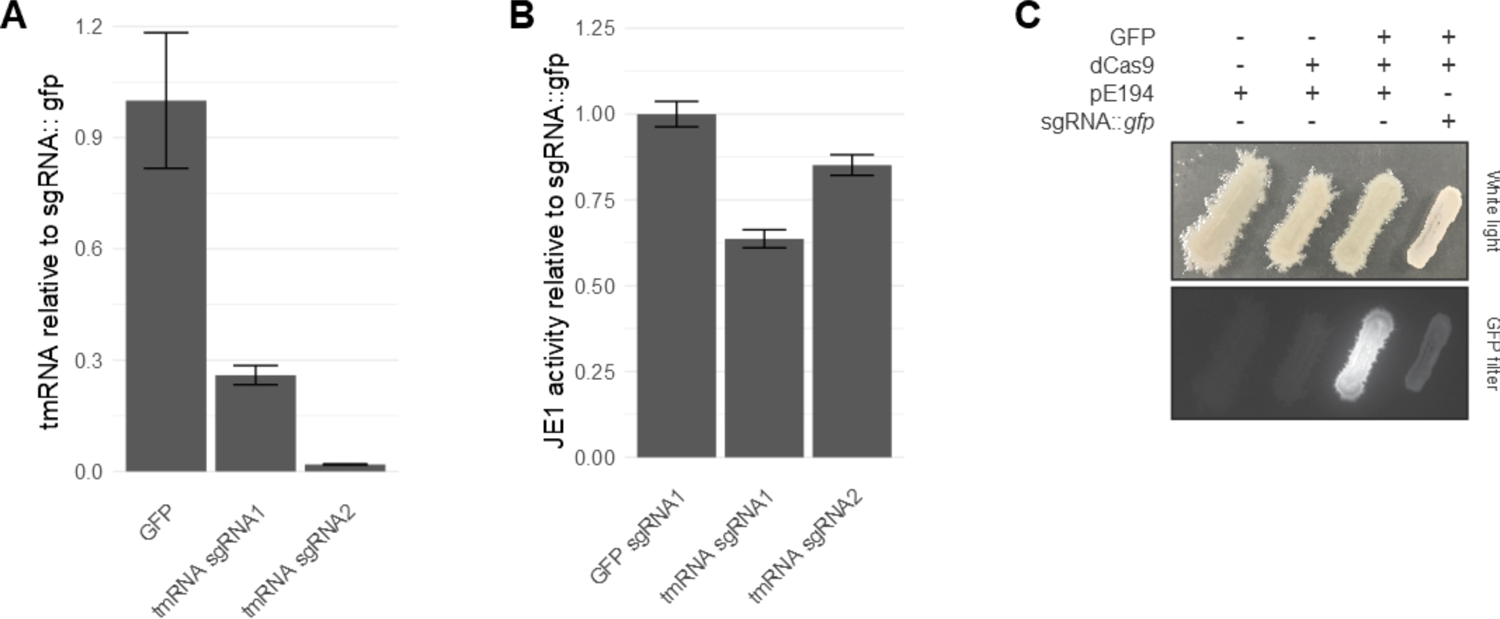
CRISPR-dCas9 functionality. (A) Quantitative RT-PCR of tmRNA in strains expressing sgRNA::*gfp* or sgRNA::*ssrA* normalized to sgRNA::*gfp* (n=3, error bars depict standard error of the mean). RNA was extracted from exponentially growing flask cultures (see methods sections “RNA purification” and “Quantitative RT-PCR”). (B) JE1 relative activity in deep-well plate fermentation strains expressing sgRNA::*gfp* or sgRNA::*ssrA* normalized to sgRNA::*gfp* (n=3, error bars depict standard deviation). (C) GFP fluorescence in strains expressing GFP, dCas9, empty plasmid (pE194), or sgRNA::*gfp*.

Each of the 53 genomic candidate regions was knocked down with 2 sgRNAs (in separate strains, as done above for tmRNA, sequences are in the Table of Supplementary file 3) and the effect on the yield of amylase was assessed with a fluorescent marker (Fig. 6 A+B). In these experiments, the introduction of the candidate targeting sgRNAs with few exceptions consistently resulted in 15.4% reduction in fluorescence compared to the control sgRNA targeted against *gfp*. This observed difference in fluorescence might be due to a global impact on transcription when dCas9 is chromosomally bound. Indeed, all sgRNAs targeting candidates direct dCas9 to the chromosome whereas the reference strain targets *gfp* localized on a plasmid. Further, this generally reduced activity pattern is consistent with the assumption that the knockdown of most candidates did not result in substantial changes in enzyme yield (despite the prior differential expression analysis and selection effort). We therefore consider the difference of activities relative to a median activity baseline to represent the changes in yield due to a specific sgRNA experiment. In fact, almost all activities are within the interquartile range (Fig. 6 A+B), underlining that most of the tested sgRNAs did not affect the yield. Among the 21 sgRNAs targeted against the genomic regions of the 15 putative ncRNA transcripts and the 6 known RNAs (Fig. 6A), we found that one of the sgRNAs targeted towards a potential novel UTR for ybfI reduced yield while one of the sgRNA targeted against *scr,* which encodes the signal recognition particle (SRP) RNA increased yield. Considering that *scr* is an essential gene, as it is involved in the recognition of signal peptides and is required for protein secretion and membrane integration, this finding was very surprising because we anticipated substantial changes in yields. Therefore, we retested the sgRNAs targeted against *scr* together with a sgRNA targeting *je1zyn* (Fig. S4.1A) in order to confirm that CRISPRi remained functional in these strains. However, we found that the viability of sgRNA::*scr* could indicate a loss of functionality of the CRISPRi system in this strain (Fig. S4.1B, see discussion for further elucidation).

**Figure 5.**
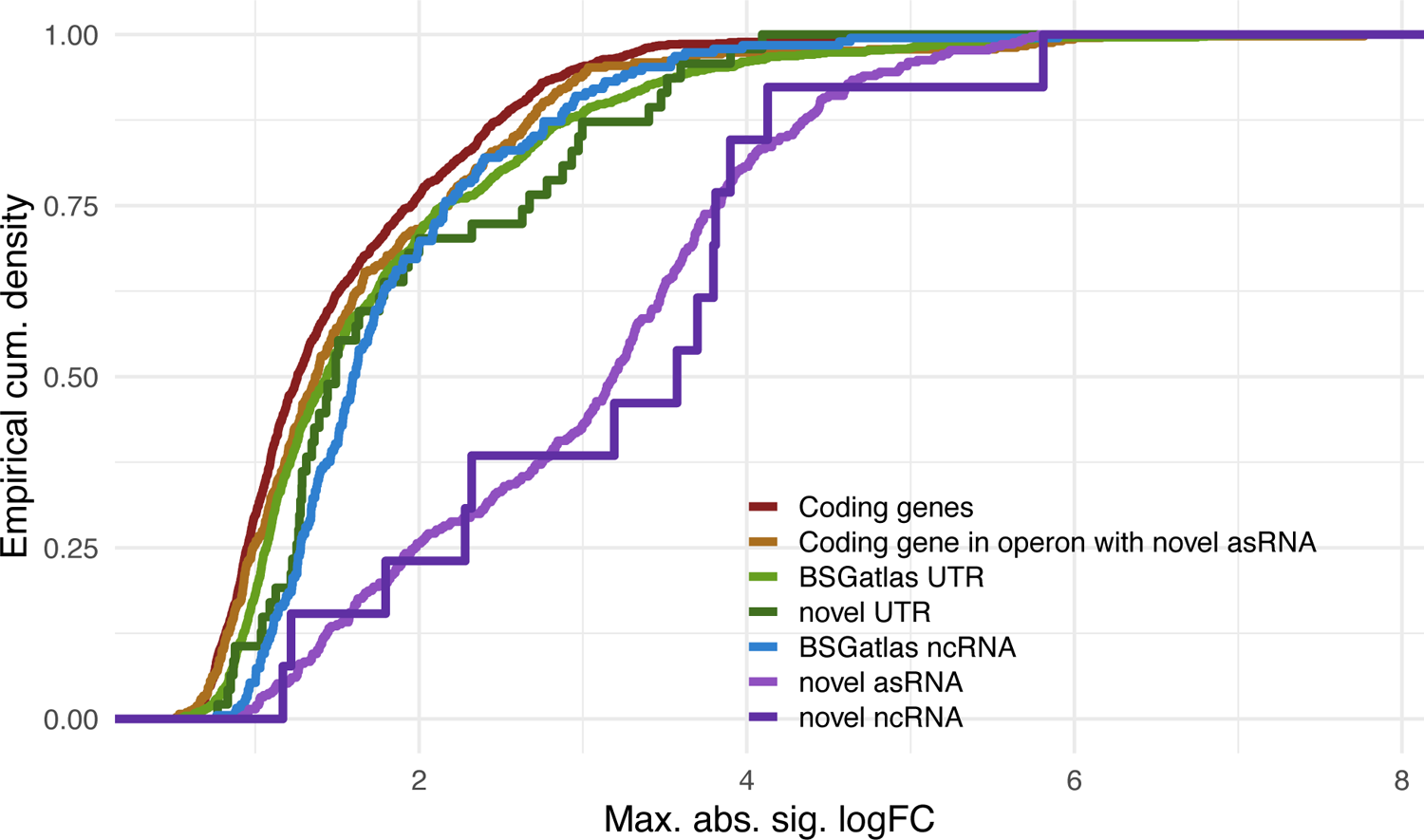
Maximum logFCs. For annotations with differentially expression, the cumulative density of the maximal observed logFC at each statistically significant pairwise comparison (see “Differential gene expression and pathway analysis”) is shown (x-axis). Coding genes are shown separately from coding genes that are located in an operon located antisense to a novel predicted asRNA.

**Figure 6.**
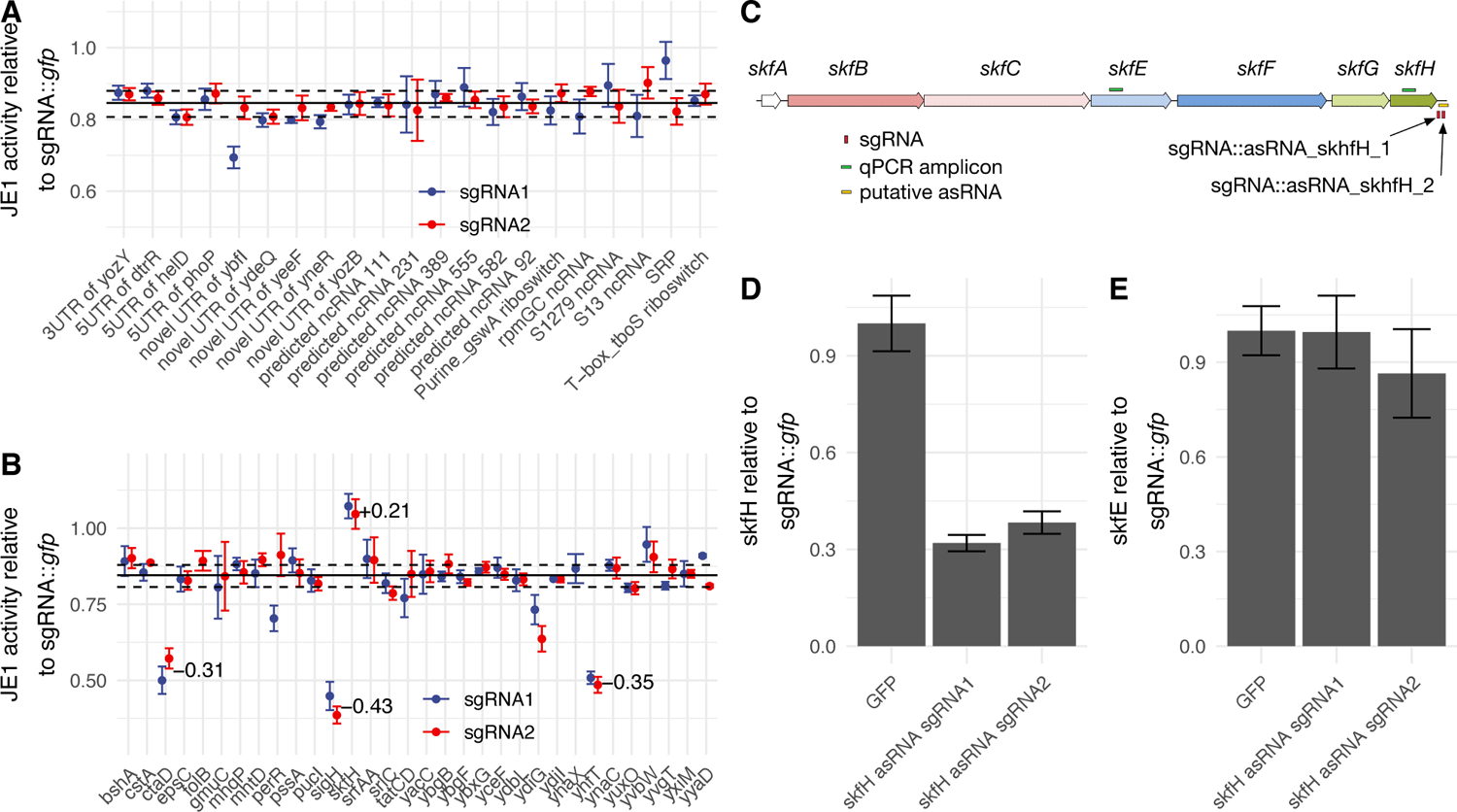
Non-coding RNA interference for impact on yield. (A) JE1 relative activity in deep-well plate fermentation of strains expressing sgRNA against ncRNA candidates. Each candidate was tested with two sgRNAs. Samples were normalized to JE1 activity of a strain expressing sgRNA::GFP (n=3, error bars depict standard deviation). Based on the observation of a consistent relative JE1 activity combined with the overall CRISPRi impact (see results), the median activity (solid black line) of all guides (including second panel B) indicates a baseline for retrieving the changes in yield. The dashed lines indicate the upper and lower interquartile range. (B) as in (A) but for 31 asRNA candidates. Data for folB sgRNA1 and yhaX sgRNA2 is not shown since these strains did not grow in liquid cultures. The average differences in yield relative to the median for the four targets ctaD, skfH, yhfT, and sigH are highlighted. (C) The *skfA-H* operon showing amplicons in qRT-PCR (green boxes), sgRNAs (red boxes) and putative asRNA (yellow box). (D) Quantitative RT-PCR of skfH coding region in strains expressing sgRNA::*gfp*, sgRNA::*skfH* sgRNA1 or sgRNA::*skfH* sgRNA2 normalized to sgRNA::*gfp* (n=3, error bars depict standard error of the mean). RNA was extracted from exponentially growing flask cultures (see methods sections “RNA purification” and “Quantitative RT-PCR”). (D) As in (C) but qRT-PCR of skfE coding region.

Among the 32 CRISPRi knockdowns targeting the genomic regions candidates selected from the list of differentially expressed predicted novel asRNAs, we find 4 regions with substantial changes in yield (Fig. 6B). One lead to an increase in yield while the other lead to a decrease. The yield increased by 21% when targeting the 3’ UTR of *skfH* and its overlapping predicted asRNA, which is activated in the early onset of nutrient stress to mediate cannibalism (Gonzalez-Pastor, 2003). Decreases in yield were observed when targeting the regions of genes or their UTRs with overlapping predicted asRNA for the cytochrome *c* oxidase subunit 1 (*ctaD*), the putative long-chain fatty-acid-CoA ligase *yhfT* (Kraas et al., 2010), and the sigma factor *sigH.* The yields were reduced by 31%, 35%, and 43%, respectively. The reduction of yields in these genomic regions are noteworthy in so far as the associated genes *ctaD* is an important enzyme in the respiratory chain and the pre-protein translocase subunit *secE,* which is part of the *sigH* operon and important for protein secretion (Song et al., 2015).

Transcription from the sense (relative to the coding gene) and anti-sense stand and the corresponding regulation can be quite subtle and complex (Waters & Storz, 2009). Thus, other genes in the *skfABCEFGH* operon are potentially affected by the 3’ UTR genomic region knockdown. In order to evaluate to what extend the genes in the operon are affected in transcription levels and to what extend the yield change is due to the knockdown of the genomic candidate region, we conducted qRT-PCR experiments with amplicons designed for both the *skfH* coding sequence and the *skfE* gene located further upstream roughly in the center of the operon (Fig. 6C). RNA was extracted for exponentially growing flask cultures (see methods “Quantitative RT-PCR”). We found that the *skfH* amplicon signal was reduced 60-70% compared to a strain containing a sgRNA targeting *gfp* (Fig. 6D). In contrast, the signal obtained for the qRT-PCR amplicon targeting the *skfE* coding region was not affected by the two sgRNAs targeted against the asRNA (Fig. 6E). This suggest that the increase in JE1 yield observed for these sgRNAs was not dependent the blocking of expression for the entire *skfABCEFGH* operon.

## Discussion

In this study, we screened for impact on *α*-amylase yield in *B. subtilis.* The main goal was to find genomic candidate regions which when targeted with CRISPRi knockdowns could result in yield changes. Given that non-coding RNAs in general are understudied in this context and given that ncRNAs have proportionally much larger logFCs (Fig. 5), we focused on ncRNAs and identify four such regions.

Our efforts involved increasing the pool of candidates to screen for, and we predicted transcribed regions. We found the antisense and other ncRNA to be most differentially regulated (in logFC values) compared to, for example, known protein coding genes (Fig 5). The RNA-seq data from fed-batch fermentation samples of two different production strains sampled at two different time points, resulted in 625 novel ncRNA candidates. Here, the transcribed regions were identified by the ANNOgesic tool which utilizes the RNA-seq read coverages (Yu et al., 2018). Known gene annotations from the BSGatlas were used to both benchmark the prediction of transcribed regions and extract novel transcribed regions (Geissler et al., 2021). Also, relative to these known annotations, the novel transcribed regions were classified as potential 106 UTR, 444 asRNA, 15 sRNA and 13 expressed regions with ambiguous annotation. We argue that the high sequencing depth applied in our study, combined with the fed-batch fermentation condition allowed us to detect these novel ncRNA candidates. Notably, 69.6% of the novel ncRNA candidates have relatively low expression levels (DESeq2 base mean below 50) and further experimentation will therefore be needed to assess their molecular characterization. Detecting the specific types (5’, 3’, internal UTR) would require additional information about transcription start and termination sites (Geissler et al., 2021; Nicolas et al., 2012). An investigation with paired-end sequencing might allow for elucidating the association (Leonard et al., 2019), yet the tradeoff in paired-end insert size would have reduced sensitivity of short transcripts. Therefore, the predicted UTRs were not further classified into the sub-types. Despite the choice of single-end sequencing in this dataset, the prediction of the 13 potential ncRNA cases should be sufficiently reliable because the prediction cut-off is larger than for 99% of all known UTRs (Geissler et al., 2021).

We determined differential expression for all genes inclusive predicted novel ncRNA using a stage-wise testing procedure (Fig. S3.4) (Love et al., 2014; Van den Berge et al., 2017), which substantially increased sensitivity (see “Differential gene expression and pathway analysis“). A surprising finding from the expression analysis was that the codon-optimized version of the enzyme had significantly higher mRNA expression levels than the non-optimized sequence, suggesting that in addition to increased translation, increased stability of the mRNA contributes to the enhanced enzyme yield observed for this strain. Based on the RNA-seq data, the transcript of the recombinant enzyme was the fourth most highly expressed RNA with two ncRNAs having even higher transcription levels: The srlX/ S1532 ncRNA and the bsrA global expression regulator 6S RNA (BARRICK, 2005). We clustered the expression profiles of both ncRNAs and protein coding genes followed by an enrichment analysis in order to assign potential pathway and Gene Ontology categories to the ncRNA candidates. Due to the observed larger magnitude of logFC for novel predicted RNA (Fig. 5), we put focus on the predicted RNAs. We used this guilt-by-association annotation of the nRNAs to select a set of 53 differentially expressed candidate ncRNAs for further testing of their effect on enzyme yield.

To screen the selected candidates for effect on yield, we implemented a CRISPRi knockdown system based on a dead Cas9, similar to previously described methods (Larson et al., 2013; Peters et al., 2016). We used sgRNAs targeted against tmRNA to validate the ability of the CRISPRi system to knockdown ncRNA expressions. Using qRT-PCR we observed efficient knockdown of tmRNA, suggesting that the set-up can be used for analysis of the impact of non-coding RNA on enzyme production (Fig. 4A). Moreover, sgRNAs targeted against GFP efficiently inhibited fluorescence of a GFP expressing strain, demonstrating the inhibition of protein coding genes (Fig. 4C). When we targeted the essential *scr* gene, encoding the SRP RNA, we observed inactivation of the CRISPRi system with one of the used sgRNAs (Fig. 6C). We expect a selection pressure to inactivate knockdown of essential genes such as *scr* and thus, the inactivation demonstrates the efficiency of the CRISPRi system. An alternative explanation for reduced knockdown of *scr* could be that the guide targets potential off-targets. However, a recent genome-wide scan for potential sgRNA off-targets in *B. subtilis,* using the binding energy-based CRISPRoff method (Alkan et al., 2018; Geissler et al., 2021), did not indicate likely off-targets for neither *scr* nor the 4 targets that substantially affected yields (Fig 6B) (any off-target sides have at least 4 mismatches and the binding specificity of guide RNA sequences was high, >9 up to 17) (Alkan et al., 2018; Geissler et al., 2021).

We applied the CRISPRi system for screening of the 53 genomic candidate regions, which were selected based on the analysis above. The selection criteria focused on ncRNA and operons potentially regulated by antisense RNA, because pure coding gene focuses have already been explored (Hohmann et al., 2016). For most of the sgRNAs, we consistently found that the enzyme yield was reduced by 15% compared to the GFP control sgRNA (Fig. 6 A+B). However, the knockdown of some ncRNA significantly affected enzyme yields. Most importantly, the two sgRNAs targeting a novel RNA expressed antisense to the *skfH* 3’ UTR showed a significant increase in yield. *skfH* is the last gene of the *skfABCEFGH* operon, which is transcriptionally induced by the Spo0A transcription factor in response to nutrient starvation (Gonzalez-Pastor, 2003). The *skfA* gene encodes a killing factor, which is secreted and mediates the killing of sister cells in proximity, thereby releasing nutrients to the cannibal cell. The remaining genes in the operon are potentially involved in the activation/release of SkfA (*e.g., skfB* and *skfC*) and making the cannibal cells resistant to SkfA (*skfE* and *skfF*). The functions of the *skfG* and *skfH* genes are unknown. Interestingly, the deletion of *skfA* has previously been shown to increase the biomass obtained from shake flask cultures, indicating that SkfA dependent lysis could influence fermentation yield (Wang et al., 2014).

One weakness of the CRISPRi system is the lack of strand-specific knockdown, meaning that both the transcription of sense and antisense transcripts will be blocked by the recruitment of the sgRNA-dCAS9 complex to the genomic sequences (Qi et al., 2013). Thus, the observed increase in yield could originate from knockdown of the putative asRNA or alternatively from blocking the sense transcription of *skfH* 3’ UTR. We find that the transcript levels at the *skfH* locus decreased upon sgRNA targeting of the 3’ UTR region, whereas transcript levels of the upstream gene *skfE* are unaffected (Fig. 6C and D). This is consistent with the observed effect on yield not being dependent on decreased expression of the first part of the operon containing *skfABCE* and therefore also consistent with the mechanism being different than the reduced lysis observed for the deletion of *skfA* (Wang et al., 2014). The qRT-PCR assay is not strand specific, meaning that the decreased expression could be due to the inhibition of the predicted asRNA or the stalled RNAP in the 3’ UTR of the operon, particularly for the expression of *skfFGH* genes. However, stalled RNAP was recently shown to be resolved by the *Bacillus* 5’ exonuclease RNase J1 (Šiková et al., 2020), which should degrade the entire transcript, which we do not observed in the qRT-PCR assay. Therefore, it could be possible that the predicted asRNA play a role in the observed increase in enzyme yield by unknown mechanism that will be the subject of further research beyond this study.

We furthermore identified several sgRNAs having a negative effect on yield. For two specific sgRNAs; sgRNA1 against *folB* asRNA and sgRNA2 against *yhaX* asRNA, we found that the strains did not grow in liquid culture. The function of yhaX is unknown, while FolB is an essential gene in Bacillus which is involved in folate biosynthesis. The observed phenotype of sgRNA1 against folB is consistent with increased requirement for tetrahydrofolate biosynthesis pathway products in liquid culture (Koo et al., 2017). In addition, we find a reduced JE1 enzyme yield from the two sgRNAs targeted against an asRNA for *ctaD*, which encodes the catalytic subunit of the cytochrome c oxidase complex that is essential for oxidative phosphorylation. The sgRNAs also block transcription of the sense strand. Thus, the reduction is plausibly also due to reduced ATP synthesis cause by the knockdown of cytochrome c oxidase complex subunit 1. Likewise, the two sgRNA targeted against an asRNA to *sigH* might affect the yields by reducing the expression of the sense *sigH-rpmGB-secE* operon. Even more so, because SecE is part of the protein secretion translocation channel, which is known to influence α-amylase expression in *Bacillus* (Mulder et al., 2013). Finally, we find that two sgRNAs targeted against *yhfT* result in a reduction in the yield and again; we hypothesize that the impact on yield stems from inhibiting expression of YhfT that is involved in the biosynthesis of the quorum-sensing molecule surfactin (Kraas et al., 2010).

Given these observed changes in yields for the *α*-amylase upon CRISPRi of genomic candidate regions, one could speculate that the targeting of the these candidates might also impact other enzymes and proteins that are secreted by the Sec pathway (Tjalsma et al., 2004); the pathway that secretes the *α*-amylase. The candidates tested in this study relate to strains under potential secretion stress due to overexpressed amylase in commercial strains (see results “RNA sequencing of bacterial enzyme production”). Further, the candidates affecting yield were associated to mechanisms that improve biomass and energy metabolism (see above). Therefore, the observed knockdowns might have affected the ability of *B. subtilis* to cope with stressful condition without directly affecting the secretion pathway. Consequentially, we hypothesize that the knockdown of the candidates might also improve yield of other Sec-pathway secreted enzymes, if their overexpression results in similar metabolic conditions.

In conclusion, we established a pipeline for the identification of putative RNA candidates that pointed to genomic candidate regions, which when interrupted by CRISPRi affect the yield of *α*-amylase production from an RNA-seq analysis of protein coding and ncRNA genes during fermentation. Further, we assessed the yield impact of the candidates with a CRISPRi based set-up. Using this strategy, we found that two sgRNAs targeted against the genomic region of a predicted ncRNA expressed antisense to the 3’ UTR of the *skfA-H* operon both led to ∼ 21% increase in yield.

## Author contributions

ASG and EGT wrote the initial draft. ASG, CA, and FA computed the novel RNA, differential expression, and pathway analysis. LDP and AOF extracted the RNA. LDP prepared the sequencing libraries. TBK conducted the CRISPR screen. AOF performed the qRT-PCR. AEB prepared the parental strains and TBK prepared the CRISPRi strains. TBK, AEB, JV and MDB coordinated the biotechnological interpretation of the results. AOF, JV, and JG revised the manuscript. CH, JV, and JG supervised the project. JG was the main coordinator of this project. All authors read and approved the final version of the manuscript.

## Acknowledgments

The authors would like to thank Lars Juhl Jensen and Nadezhda Doncheva for their feedback on methodology. We thank Anette Holtmann for sgRNA cloning assistance.

