## Supplementary File 1 for "CRISPRi screen for enhancing heterologous *α*-amylase yield in *Bacillus subtilis*"

### Additional File 1
### supporting online material

#### S1 Deep RNA-sequencing

The library was reverse transcribed with random priming position with the adapter 5’-AGACGTGTGCTCTTCCGATCTNNNNNNNNNNNNNNN. Afterward, the adapter 5’-PHO-NNNNNNNAGATCGGAAGAGCGTCGTGTAGGGAAAGAGTGT was ligated to the 3’ end of the cDNA. The cDNA library was PCR amplified with the primers 5’-AATGATACGGCGACCACCGAGATCTACACTCTTTCCCTACACGACGCT and the multiplexing adapter 5’-CAAGCAGAAGACGGCATACGAGAT*iiiiii*GTGACTGGAGTTCAGACGTGTGCTCTTCCGATCT. The resulting reads have the sequence structure
5’-AATGATACGGCGACCACCGAGATCTACACTCTTTCCCTACACGACGCTCTTCCGATCT + 7 random PCR barcode + payload + AGATCGGAAGAGCACACGTCTGAACTCCAGTCAC*jjjjjj*ATCTCGTATGCCGTCTTCTGCTTG-3’, where *jjjjjj* is the reverse complement of the multiplexing index *iiiiii* (Table S1.1).

**Table S1.1**. Multiplexing indices and information associated with the RNAseq libraries. The experiments per day and strain were run with duplicate samples.

| Strain | Sample | Day | Index |
|---|---|---|---|
| ***je1 + prsA*** | MIX1441 | 1 | ATCACG |
| ***je1 + prsA*** | MIX1442 | 1 | CGATGT |
| ***je1zyn + prsA*** | MIX1443 | 1 | TTAGGC |
| ***je1zyn + prsA*** | MIX1444 | 1 | TGACCA |
| ***je1 + prsA*** | MIX1441 | 3 | ACAGTG |
| ***je1 + prsA*** | MIX1442 | 3 | GCCAAT |
| ***je1zyn + prsA*** | MIX1443 | 3 | CAGATC |
| ***je1zyn + prsA*** | MIX1444 | 3 | ACTTGA |

**Table S1.2.** The number of reads before and after filtering, including mapping statistics. Each column's percent points are relative to the number of reads in the column directly on the next column to the left. MIX1441 and MIX1442 are of substrain *prsA+je1*, while MIX1443 and MIX1444 are of substrain *prsA+je1zyn*.

| Sample | Day | Raw reads | Cleaned reads | Mapped reads | Uniquely mapping reads |
|---|---|---|---|---|---|
| **MIX1441** | 1 | 24,236,309 | 18,881,911 (77.91%) | 18,812,375 (99.63%) | 18,553,361 (98.62%) |
| **MIX1442** | 1 | 22,691,223 | 16,734,508 (73.75%) | 16,665,005 (99.58%) | 16,562,262 (99.38%) |
| **MIX1443** | 1 | 24,069,205 | 18,305,729 (76.05%) | 18,173,946 (99.28%) | 17,987,875 (98.98%) |
| **MIX1444** | 1 | 27,982,513 | 18,770,284 (67.08%) | 18,622,749 (99.21%) | 18,467,752 (99.17%) |
| **MIX1441** | 3 | 29,061,221 | 15,684,975 (53.97%) | 15,567,946 (99.25%) | 15,495,214 (99.53%) |
| **MIX1442** | 3 | 9,186,714 | 5,342,049 (58.15%) | 5,307,152 (99.35%) | 5,259,570 (99.1%) |
| **MIX1443** | 3 | 21,027,577 | 17,665,779 (84.01%) | 17,560,343 (99.4%) | 17,472,234 (99.5%) |
| **MIX1444** | 3 | 22,425,410 | 16,076,356 (71.69%) | 15,981,246 (99.41%) | 15,919,519 (99.61%) |

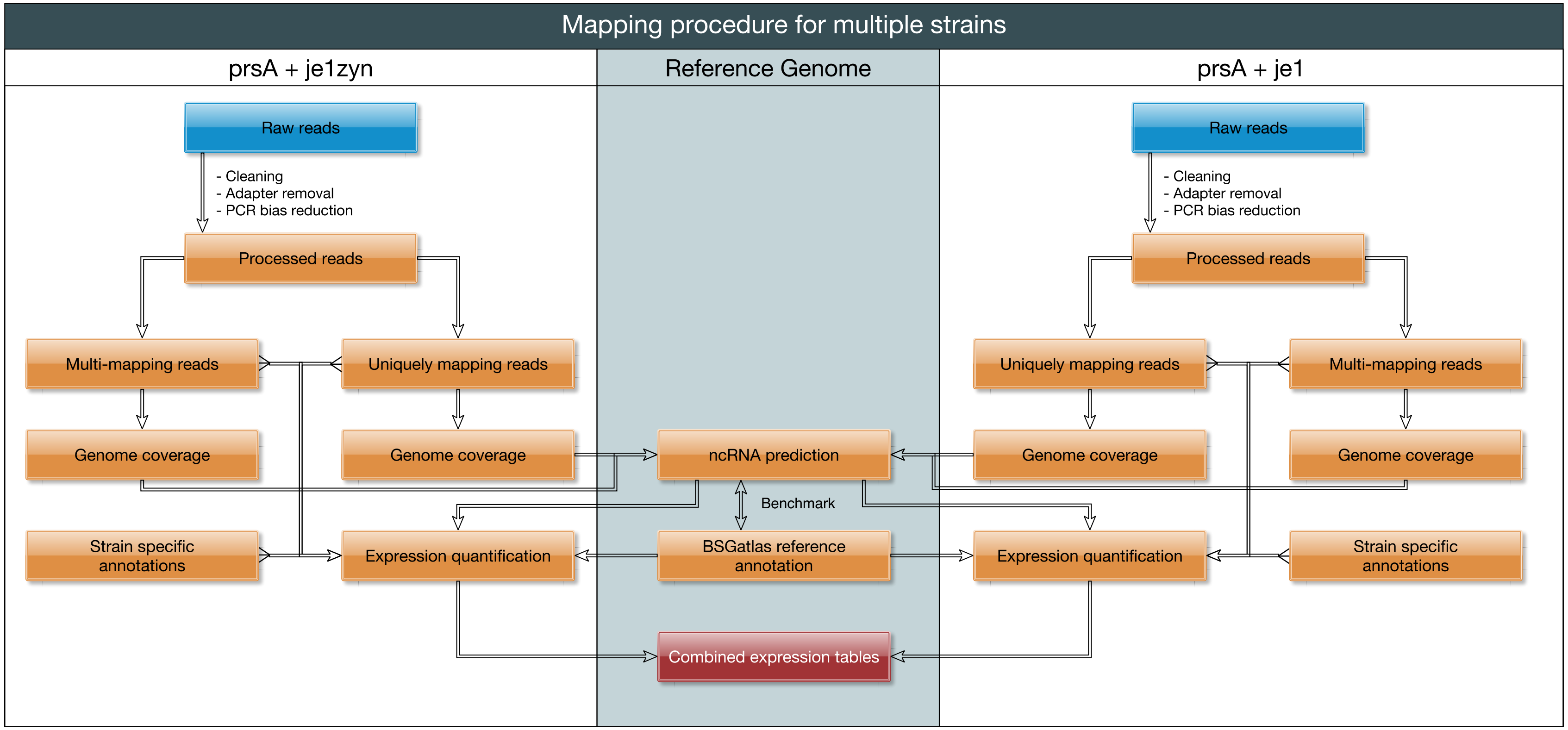


**Figure S1.3**. Analysis Workflow. The RNAseq libraries (blue) were similarly pre-processed and mapped against the genome sequences of the two respective strains (orange-colored steps). Genome coverages of the individual strains were computed for multi-mapping and uniquely mapping reads and normalized relative to the respective number of reads. Coverages were lifted over to the RefSeq reference genome sequence for the prediction of novel ncRNAs. The ncRNA prediction was benchmarked against the BSGatlas annotation. The resulting predicted ncRNA positions and BSGatlas annotation were lifted to the two strains for quantification of expression together with the strain-specific genes. The two expression tables, one for each strain, were combined into a single table (joined on gene identifiers) which was then used as input (red) for the subsequent expression analysis steps (differential expression and pathway analysis).

#### S2 Benchmark and prediction of novel ncRNA

The two main parameters that influenced the computation of predicted transcribed regions (PTR) were the height cut-off in expression coverage and the number of replicates a prediction should be observed (Fig. 2; (Yu, Vogel, and Förstner 2018)). The tool also tolerates regions of a few consecutive base-pairs (tool default: 5 bp) to be below the height cut-off. The tool further automatically increases the tolerance for regions that are within already annotated transcripts. Here, we benchmark how the parameters affect recall of known reference transcript annotations from the BSGatlas v.1.1 (Geissler et al. 2021), which provides 4,879 reference transcript annotations, of which only 362 annotations were for genes with unknown transcripts or operon. In total, 4,403 reference annotations were coding transcripts, 375 non-coding transcripts, and 101 more complex transcripts, such as those containing both coding sequences and small RNA structures (sRNAs). We tested transcript region prediction with ANNOgesic for 80 different parameter combinations (all height cut-offs 1 through 20 for our 4 replicates). For all parameter settings, the number of predictions, lengths, and overlap relationship to reference transcript annotations were examined. After the benchmark, we selected the ideal set of parameters and extracted potential novel ncRNA annotations. The details of these steps are shown here.

Lowering the height cut-off increased the number of PTR exponentially (Fig. S2.1 A). Allowing predictions to be observed in a single replicate instead of requiring them to be observed in all replicates led to a four-fold increase in the number of predictions (Fig. S2.1 A). Given that stronger expression signals and multiple observations coincide with confidence in predictions, a more conservative set of predictions are, as observed, characterized by a high cut-off value and required each prediction to be observed in all samples. More relaxed settings had lower cut-off values and fewer observations accordingly. When relaxing the settings, the increase in predictions was particularly pronounced for short PTR (1,000 bp) at lower height cut-offs (≤5; Fig. S2.1 B). Also, the distribution of maximally observed per sample averaged coverage—a measure for overall expression—drastically decreased for that value (Fig. S2.1 C).

We investigated how relaxing the prediction parameter impacts the recall of known transcripts. The prediction parameters with overlaps between known transcripts and predictions in the absolute number of base pairs and relative to the prediction/transcript lengths were checked. Furthermore, the numbers of predictions that do not overlap with known transcripts, and those predictions that also had a low expression level, were determined. The latter indicated at what parameters the increase in sensitivity becomes disproportional to the noisy, putatively erroneous predictions. The overall distribution of overlaps between the PTR and the reference set (Fig. S2.2) showed that the strict prediction parameter, PTR predominately overlapped the first half (closer to 5' end) of the reference annotations. In contrast, at the relaxed settings, the predictions start much further downstream and end much further upstream from the reference annotations–at least twice the annotations' length in both directions. A balance between those two extremes, at which the PTR were more or less equal to the references, was observed for height cut-offs between three and five. More lax predictions (low cut-off with few or only one observation) overlapped almost all coding and around 80% of the non-coding annotations. (Fig. S2.1 D). However, requiring a minimal relative overlap brought these ratios down to 50% in both cases. The proportion of annotations that were covered decreased the most for minimal relative overlaps above 75%. Enforcing stricter settings lowered the overall proportion, although this decrease in stricter settings was less pronounced for height cut-offs above five. Inspection of what number of PTR relative to the total number of predictions were involved in these overlaps showed that, for cut-offs above five, at least 75% of PTR were relevant. In contrast, for a cut-off of three only half the predictions and the lowest cut-off (one) less than a quarter of predictions were used (Fig. S2.1 E). The remaining PTR were almost all short (1,000 bp) and with low coverage (Figs. S2.1 F-G). These observations were similar for both coverage versions. Such that gene copies were considered, this benchmark was repeated for both coverages for all mapping reads, which included multi-mapping reads, and the separate profile for only uniquely mapping reads. The overall trends were the same. Given these observations, we continued with the predicted transcripts resulting from the parameters height-cut offs five and observation by at least two replicates.

The length of gaps between known transcript annotations (Fig. S2.3 A) showed that NTRs without antisense overlaps (that is, without overlap by a reverse complementary annotation) could have lengths ranging from 100 bp up to 1,000 bp. However, when also considering potential antisense overlaps, a bi-modal distribution was revealed with two modes of short gaps (<300 bp) and longer gaps that were up to 100,000 bp. Thus, there was potential for novel ncRNAs with and without antisense overlaps. PTRs that intersect, at least partially, with these gaps were identified and then extracted from these overlaps potential novel, not yet annotated transcribed regions (NTRs) (Fig. S3B). After removing short (≤10 bp), biologically unlikely fragments, NTRs were classified according to possible antisense overlaps and distances to neighboring known annotations. The above determining prediction parameters resulted in 2,072 PTR when using only uniquely mapping reads and 2,198 PTR when including multi-mapping reads. These two PTR sets were combined by merging any overlapping predictions. Afterward, potentially novel transcribed regions (NTR) were extracted as described above, resulting in a set of 1,103 NTR. The bulk of the NTR (957) was shared by the prediction from uniquely-/multi-mapping coverages, 125 were only found with multi-mapping reads, and 21 only by uniquely mapping reads. The lengths of the novel regions were mostly 69 bp up to 1127 bp long (full distribution, Fig. S2.3 B). Here, novel regions were selected by the intersection with annotation gaps, which resulted in 14.3% of very short predictions (≤ 10 bp). These novel regions are too noise because they cannot have evidence due to the read lengths (see Library preparation and sequencing). 25.2% slightly longer but still short predictions (11–50 bp) that might be covered by reads are biologically unlikely, because >96% of asRNA and sRNA and 50% of the UTRs annotated in the BSGatlas are longer. Therefore, we removed 428 short fragments (< 50 bp). The remaining NTR were classified as potential asRNA if they were overlapped with a reference transcript (asRNA to ncRNA exist in *B. subtilis, e.g.,* in the bsrE/SR5 system (Meißner, Jahn, and Brantl 2016)) in at least 90% of their length (Fig. S2.3 C). The inclusion of multi-mapping reads led to some NTR having overlaps with rRNA. Thus, we further distinguished NTR into 444 asRNA and 97 ncRNA antisense to rRNA. Interestingly, putative asRNA had a disproportionally large distance to their nearest same strand neighbor compared to the not-yet-classified NTR (Fig M2.4). The distances to the closest neighbor for the remaining NTRs were < 12,000 bp, although the bulk had an annotation closeby (100 bp). These distance thresholds were considered to classify the remaining NTR as either a UTR (106) for a closeby gene or a putative novel ncRNA (15) (Fig. S2.3 D). The putative UTR regions were not further distinguished into 5', 3', and internal UTR (see explanation in "Prediction of novel ncRNA transcripts" results section). There were only 13 cases with distances in-between both values, considered unclear/inconclusive. There was no apparent dependency between this classification of an NTR and its length (Fig. S2.3 E). Overall, a total of 675 putative novel ncRNA candidates were identified in our study.

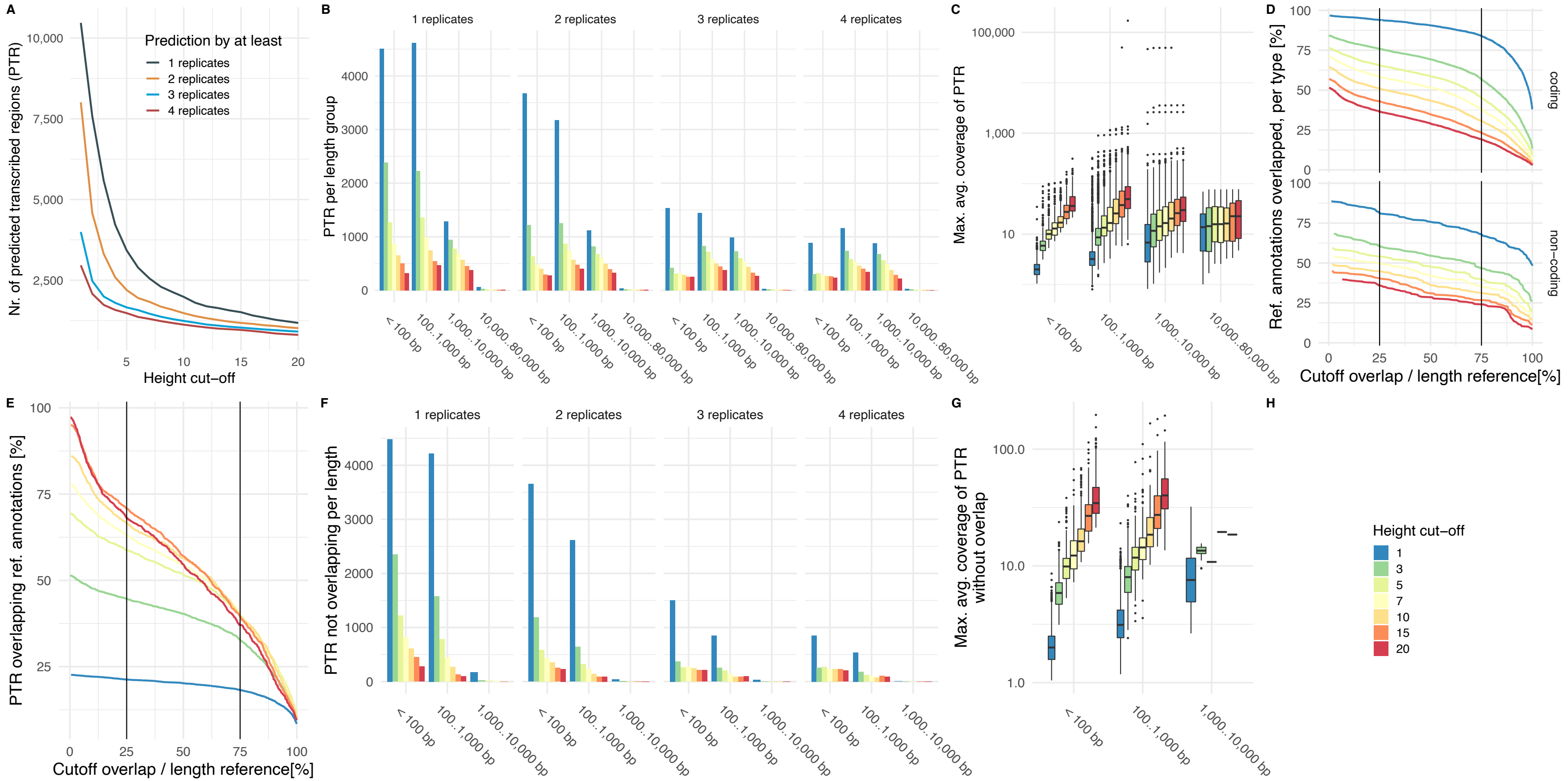


**Figure S2.1.** Summary of the benchmark for predicted transcribed regions (PTR). (A) Total number of PTRs dependent on height cut-off. (B) Length of PTR relative per type and replicate parameter. (C) Distribution of maximal per sample averaged coverage of each PTR. (D) Proportion of coding and non-coding reference annotations covered by PTR depending on the minimal required overlap. (E) The proportion of all PTR predictions that are used for the overlaps shown in D (F) Length distribution for those PTR that are not involved in a relative overlap of at least 25%. (G) Similar to C, but only for the PTR shown in F. (H) Color legend of the height cut-off used in plots B through G.

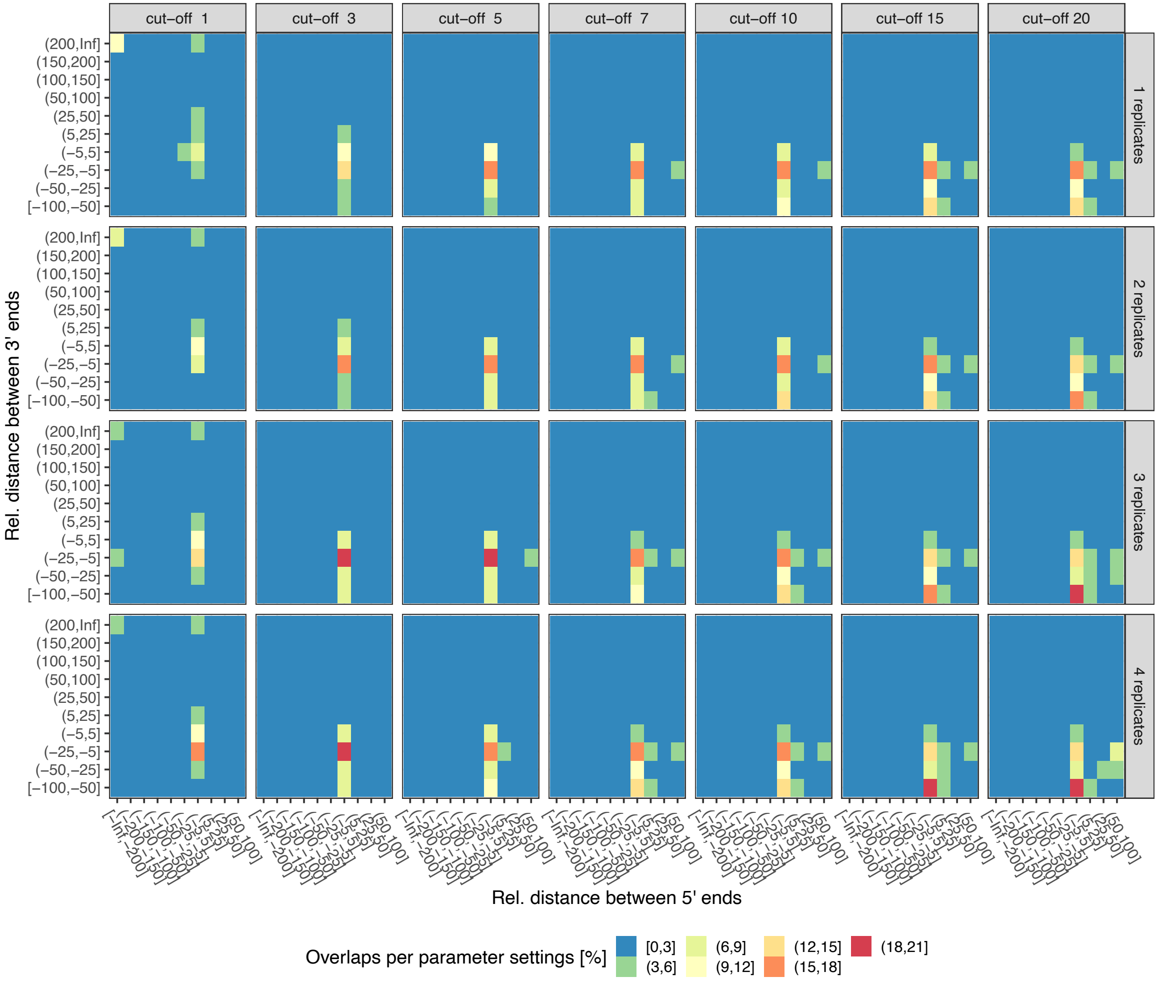


**Figure S2.2.** Large scale comparison of 5' and 3' end distances of predicted relative to reference transcript annotations. For each PTR, the distance [bp] of the 5' and 3' ends relative to the same strand overlapping reference annotations were computed as shown in Figure 2. For each parameter settings with which PTR were computed (columns: height cut-off, rows: required number of observations), the distances were counted and bins and normalized by the number of overlaps observed for the parameter setting.

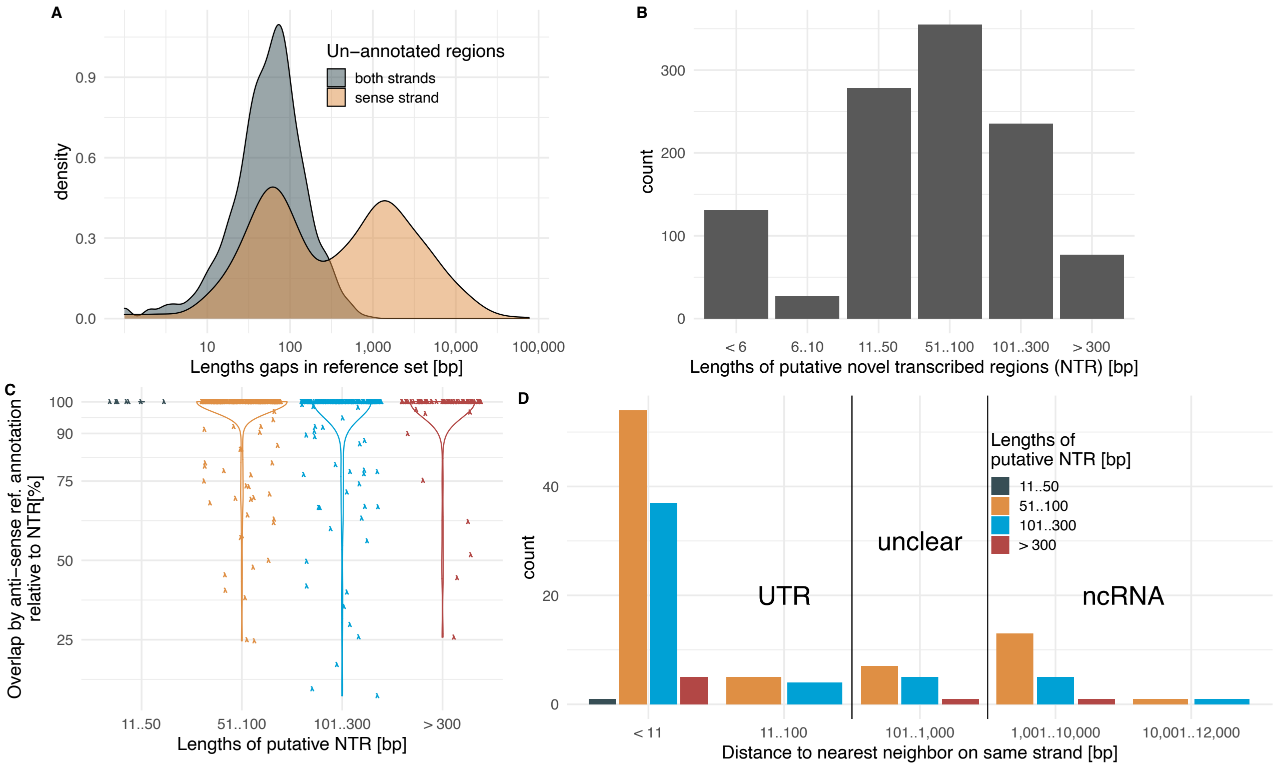


**Figure S2.3.** (A) Length distribution for gaps in annotations. (B) Length distributions of NTRs. (C) Violin plot of antisense coverage for NTRs. NTRs without any overlaps are omitted. (D) Distances to closest same strand neighbor for NTRs with antisense annotation coverage below 90% of their lengths. NTRs close to annotations were classified as potential UTRs and NTRs with larger distances as potential independent ncRNAs. The distances of 13 cases were inconclusive to classify them as UTR or ncRNA (see S2).

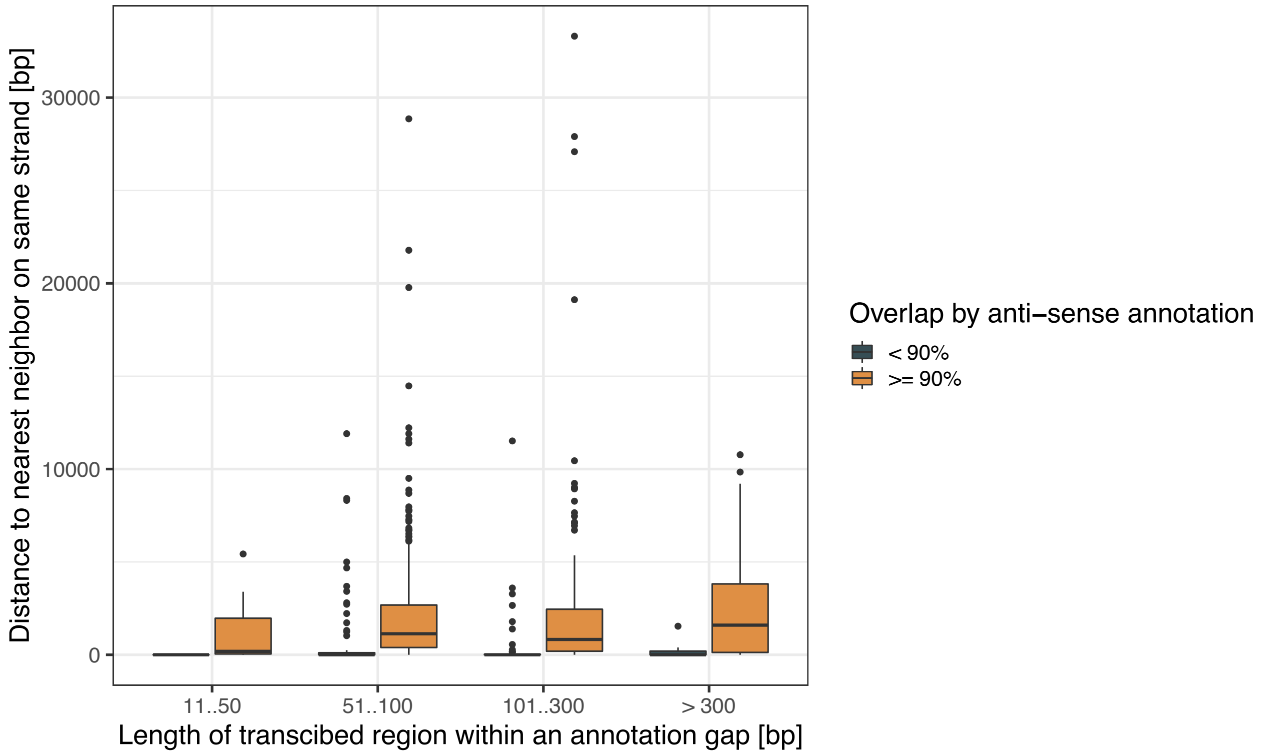


**Figure S2.4**. Illustration of distances [bp] to nearest same stand neighbor of NTR separated by the length of the NTR (x-axis) and if the NTR had an antisense overlap of at least 90% of its length or not (box color).

(A)

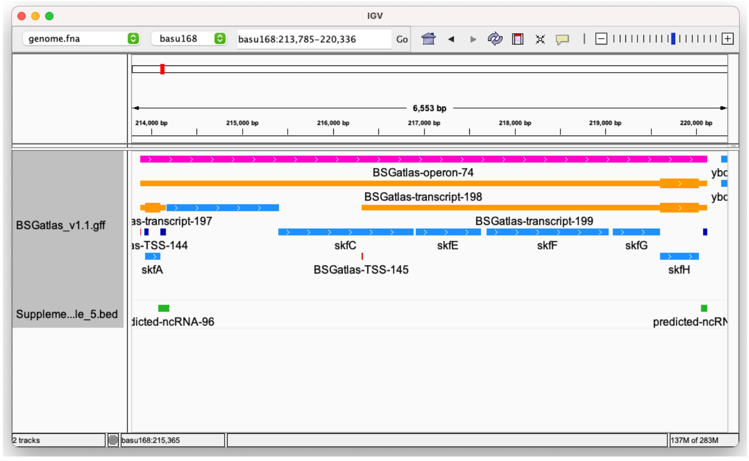


(B)

| PUTATIVE CLASSIFICATION | COLOR |
|---|---|
| Anti-sense to rRNA genes | Purple |
| UTR | Orange |
| Anti-sense or potential sRNA | Green |
| Unclear | Red |

**Figure S2.5.** Visualization of predicted ncRNA. (A) Screenshot of the integrated genome browser software (IGV) that shows the annotation of the skfABCEFGH operon in the BSGatlas next (available under https://rth.dk/resources/bsgatlas/download.php and https://doi.org/10.5281/zenodo.4305872) next a track of the predicted ncRNAs from this study (Supplementary file 5). (B) Overview over the colors used to highlight the putative classification of the predicted ncRNA (see S2). Predictions located on the reverse strand are shown in slightly darker coloration.

#### S3 Stage-wise differential expression analysis

The stage-wise paradigm is split into a screening and a confirmation stage (Van den Berge et al. 2017). This screening step is a single hypothesis test that is logically required by the confirmation step, and thus serves as an omnibus to bundle multiple hypotheses). Here, we test for differential expression at five pairwise comparisons (between strains and during fermentation within each strain, see Fig. S3.4) are applied to each genomic biotype (gene, UTR, etc.). Thus, when relating to the same feature, they share the assumption that not only is the feature expressed, but it also has a non-constant expression. Therefore, a logical preceding step first tests screens which features have changes in expression at all before confirming which of the pairwise tests are affected. When nesting the *P* adjustment of the screening and the confirmation step, an overall FDR (OFDR) control is guaranteed (Van den Berge et al. 2017). The approach is first to adjust the screening level's FDR relative to the number of all features. Afterward, the confirmation hypotheses were subjected to family-wise error correction (FWER). However, this correction is done only for those features that passed the initial screen and the adjustment is relative to the number of confirmation hypotheses scaled by the number of features that passed the screen (Van den Berge et al. 2017). Logical dependence between screening variables can be used to further improve the FWER correction's statistical power (Shaffer 1986; Van den Berge et al. 2017). For our formulated hypotheses, a trivial combinatorial analysis reveals that at least two pairwise tests must indicate differential expression once it is known that the expression is changing. Thus, two can be used as a baseline for the modified sequentially rejective multiple test procedures (Shaffer 1986). Therefore, dividing only the two smallest *P* by factor two and not changing the remaining *P* is sufficient to adjust the FWER.

The reference BSGatlas described 4,332 coding genes and 408 non-coding genes, including 104 riboswitch RNA structures and 188 non-coding RNA genes (ncRNA) that are not tRNA (86) and rRNA (30). Transcriptional information, including polycistronic transcripts, was provided for 93% of all coding and non-coding genes. That information comprises a set of 2,266 operons and 5,335 untranslated regions (UTRs), which includes 5', 3', and internal UTRs. Derived from this information, a set of 4,517 transcripts were provided in the resource. Only 362 genes are not yet annotated with a known transcript. For the expression quantification, we lifted over these annotations to the respective genome (see main methods).

**Table S3.1.** Impact of stage-wise hypothesis testing. For each coding/non-coding gene, five hypotheses of differential expression were tested (Fig. S3.4). The table compares how many genes and non-coding elements were detected as potentially differentially expressed by a direct naive combination of the pairwise tests or by a preceding screening step and afterward investigating the pairwise test in a stage-wise procedure.

| Detection per pairwise hypothesis tests | Only stage-wise testing | Only naïve approach | Both procedures |
|---|---|---|---|
| **Coding sequences** | 578 | 1 | 1,210 |
| **UTR + riboswitch** | 342 | 4 | 705 |
| **other non-coding RNA** | 40 | 0 | 401 |
| **tRNA** | 6 | 0 | 60 |
| **extra-*prsA*** | 0 | 0 | 1 |
| **JE1** | 0 | 0 | 1 |
| **total** | **966** | **5** | **2,378** |

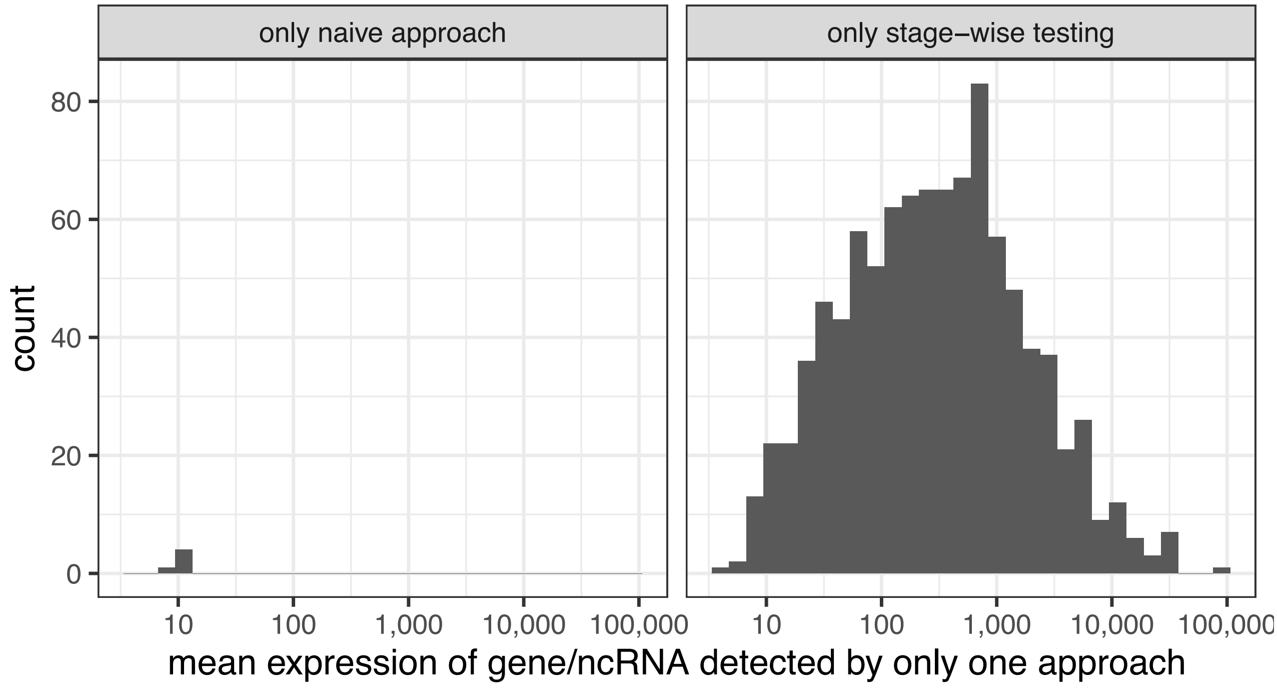


**Figure S3.2.** Mean expression strength of genes/ncRNA detected by only one either differential expression approach, the naive combination of the pairwise hypotheses tests, or the stage-wise procedure.

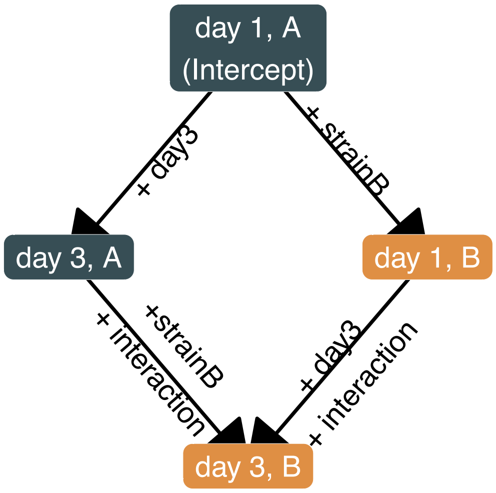


**Figure S3.3.** DESeq2's regression model for our experimental setup. In this scenario, the regression models the mean expected expression on the log2 scale for four cases. The intercept represents the base expression on day 1 for the base strain A (prsA + je1, blue nodes). The expressions for the other conditions (strain B, orange nodes) are inferred via the addition of the coefficients as shown.

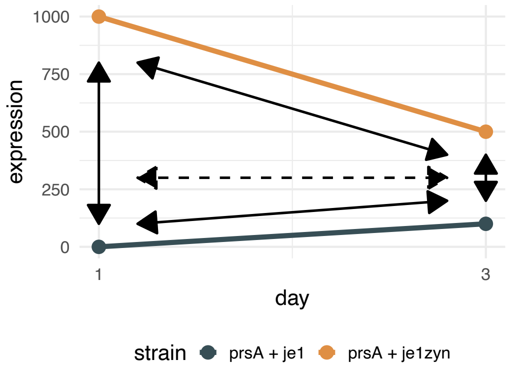


**Figure S3.4.** Pairwise, multiple hypotheses relative to the expression measurement. For each gene and non-coding element (ncRNA, UTR, etc.), four hypotheses were investigated between the strains and along time (black arrows). A fifth hypothesis (dashed arrow) tests if the change between the strains differs between the days. Given that these tests relate to the same gene profile, we first screened if the expression changes overall and only then specifying which of these pairwise tests the changed (see methods). We furthermore leveraged the screen to weigh the hypotheses in a stage-wise procedure, such that the overall FDR was guaranteed.

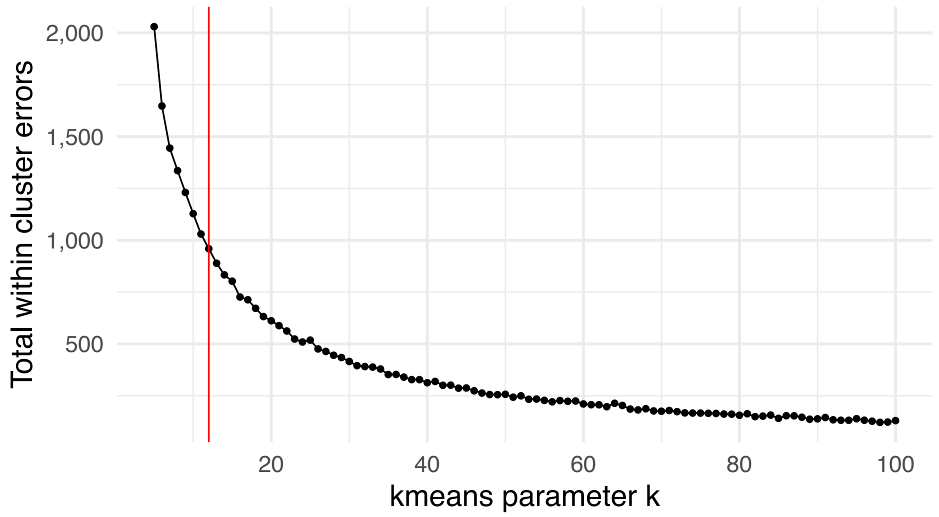


**Figure S3.5.** Defining the numbers of *k*-means clusters according to the elbow method (Thorndike 1953). Increasing this parameter reduced the total sum of within-cluster errors. For clustering input see methods.

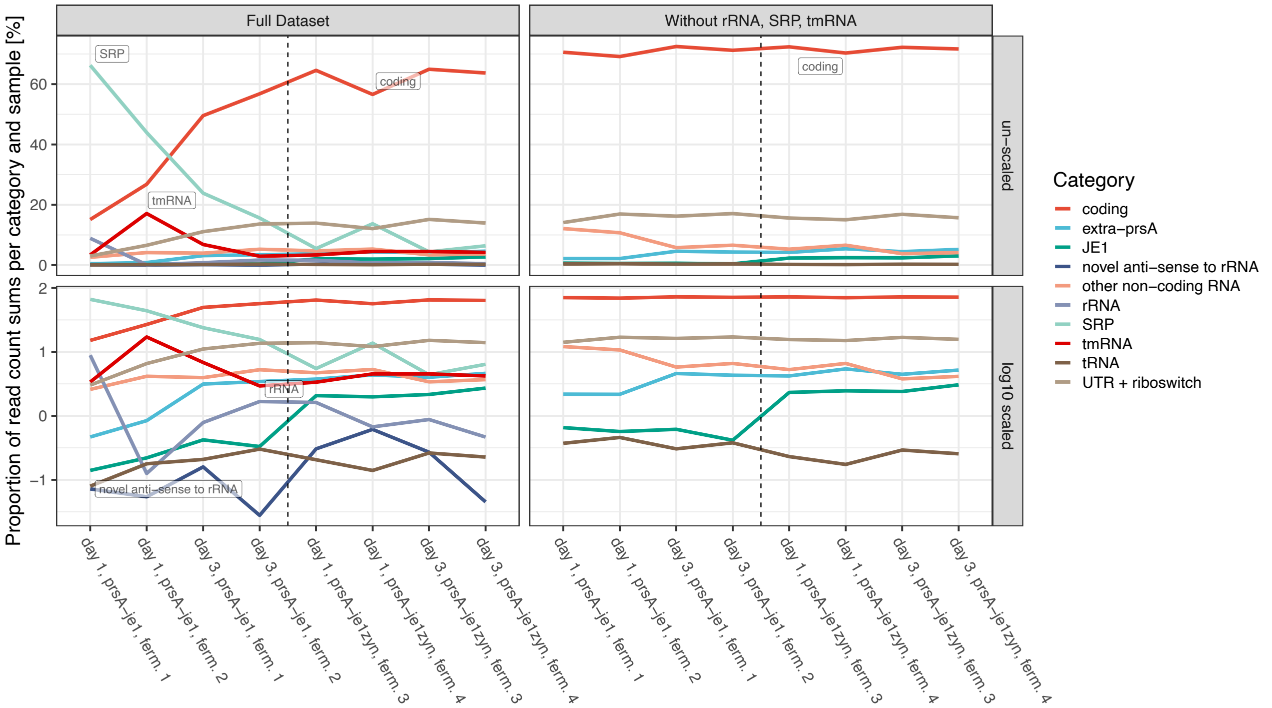


**Figure S3.6.** RNAseq library content. Shown is for each RNA-seq library the proportion of expression per gene category (sum) in relation to the total sum of expression. The proportions are shown for two cases: When considered all genes and after removal of rRNAs, tmRNAs and SRPs. The first row plots the proportions in a linear coordinate system, and the second row in a log-scale. The dashed line separates the libraries of the two strains. The shown expression quantifications included multi-mapping reads. When using only uniquely mapping reads, the expression of rRNA related genes is substantially lower, yet the overall proportion remains similar.

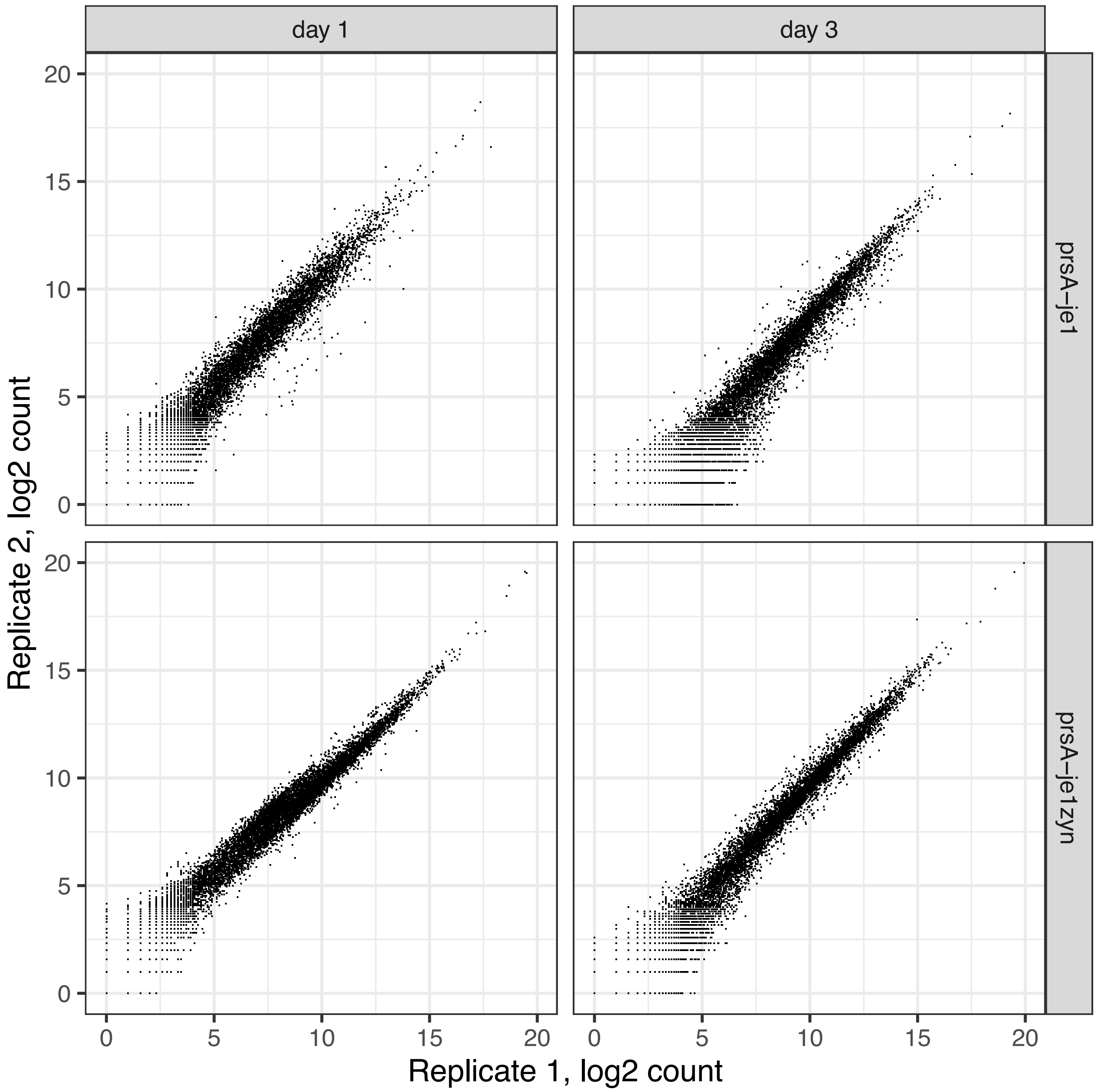


**Figure S3.7.** Scatter plot of replicates of each condition (same strain and day). Shown are the scatter plots of the raw, un-normalized expressions, excluding rRNAs, tmRNAs, and SRPs, for each pair of biological replicates.

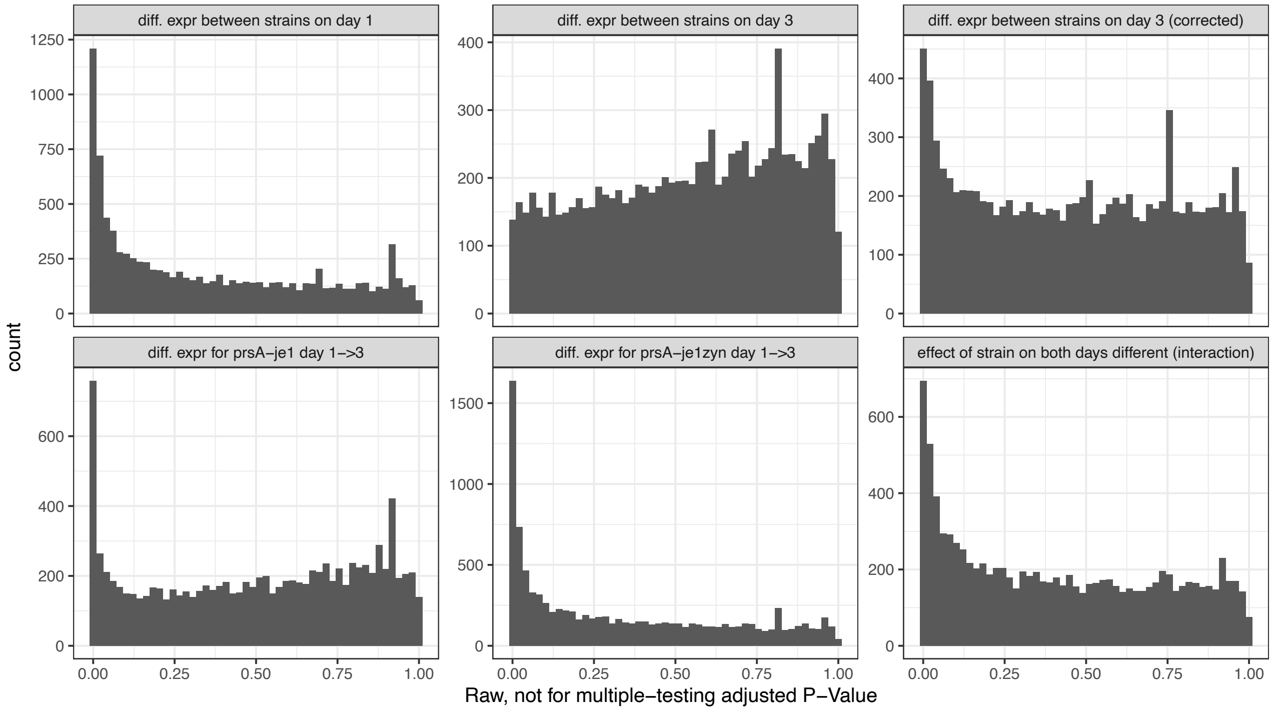


**Figure S3.8.** Un-adjusted *P*s and corrected *P*s distributions. Shown are the *P*s histograms of the un-adjusted *P*s for the 5 pairwise tests for differential expression. The distribution for differential expression between the strains on day 3 did not meet the requirement for FDR adjustment, such that these were first corrected with the *fdrtool*.

#### S4 CRISPR-dCas9 knock-down system

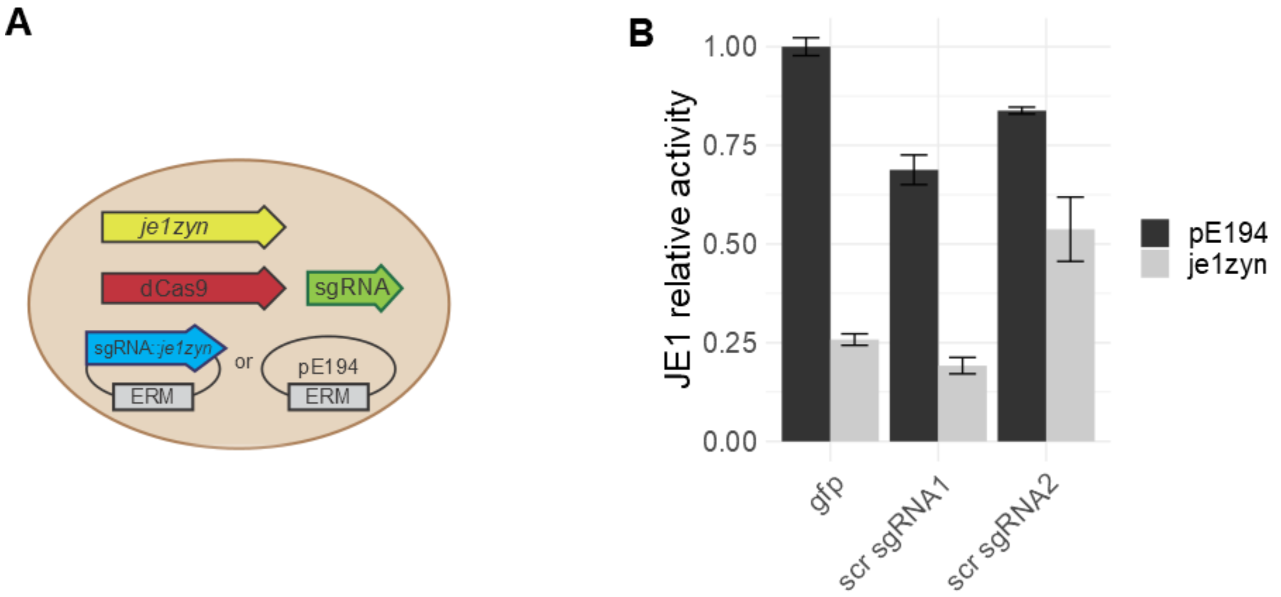


**Figure S4.1.** CRISPRi functionality test of knocking out an essential gene (A) Graphic representation of the additional verification of the dual sgRNA setup. (B) JE1 protein activity levels. Strains were either transformed with plasmid containing sgRNA*::je1zyn* (pTK0002) or negative control plasmid pE194 in strains expressing sgRNA*::gfp*, sgRNA::*scr* 1 or sgRNA::*scr* 2 (n=3, error bars depict standard deviation).
